# Determinants of CCT–motif specificity in WNK signaling and expansion of CCT-like domains

**DOI:** 10.64898/2026.04.15.718729

**Authors:** Germán Magaña-Ávila, Eréndira Rojas-Ortega, Mario Lira-Castañeda, Iván Diaz-Ortiz, Juan Bustamante, Héctor Carbajal-Contreras, Esteban Rojas-Juarez, Regina Ortega-Prado, Alejandro Marquez-Salinas, Norma Vazquez, Gerardo Gamba, María Castañeda-Bueno

**Affiliations:** Department of Nephrology and Mineral Metabolism, Instituto Nacional de Ciencias Médicas y Nutrición Salvador Zubirán, Tlalpan, Mexico City, Mexico; Facultad de Medicina, Universidad Nacional Autónoma de México, Coyoacán, Mexico City; PECEM (MD/PhD), Facultad de Medicina, Universidad Nacional Autónoma de México, Coyoacán, Ciudad de México, México; Molecular Physiology Unit, Instituto de Investigaciones Biomédicas, Universidad Nacional Autónoma de México, Tlalpan, Mexico City, Mexico

## Abstract

WNK kinases regulate ion transport and cell volume through interactions with partner proteins such as SPAK, OSR1, NRBP and TSC22D proteins. These interactions are mediated by conserved C-terminal (CCT) domains that recognize short sequence motifs, but the rules governing this recognition remain incompletely defined. Here we show that these domains can be grouped into distinct structural classes with different binding preferences for motif variants. We identify a previously unrecognized motif that mediates binding to the second CCT domain of WNKs and show that interaction specificity is determined by conserved physicochemical features, including electrostatic contacts and aromatic interactions, rather than strict sequence conservation. We further identify a similar domain in the protein FERRY3 that can bind TSC22D motifs in isolation. These findings define a framework for interaction specificity in WNK signaling and suggest that this binding mechanism may extend beyond this pathway.

## Introduction

With No lysine (K) (WNK) kinases belong to a family of four members in mammals that participate in the regulation of diverse physiological processes ^1^. They regulate transepithelial ion transport in the kidney and other tissues, modulate intraneuronal chloride concentration and thus the response to certain neurotransmitters, and participate in the cellular response to osmotic stress. Regulation by WNK kinases of the activity of the cation chloride cotransporters (CCCs) of the Slc12 family is relevant in all these processes, where inhibition or activation of cotransporter function affects membrane ion fluxes having different consequences in different scenarios, e.g., affecting ion movement through epithelia, cellular chloride concentration, or net cellular ion content, and thus, cellular volume. The best well-known substrates of WNK kinases are the Ste20-related proline-alanine-rich kinase (SPAK) and the oxidative stress response 1 (OSR1) kinase, whose phosphorylation promotes their activation ^2,3^. These kinases in turn phosphorylate the CCCs in different sites that affect their activity ^4–6^. Sodium-dependent CCCs become activated upon phosphorylation, whereas sodium-independent (and potassium-dependent) CCCs become inactivated ^1^. In addition, WNK kinases have been implicated in other processes, like cancer and T cell proliferation ^7,8^, in which the involvement of Slc12 cotransporters is less well defined.

Components of the WNK pathway have been shown to interact through a common mechanism. This mechanism involves the interaction of a globular domain named the Conserved C-Terminal (CCT) domain, because it was initially identified as a conserved region in the C-terminus of SPAK and OSR1, with a short linear motif that was initially described to contain the canonical sequence RFxV/I ^4^. The first description of this interaction mechanism was provided by Piechotta and Delpire, who identified SPAK and OSR1 as interacting partners of KCC3, NKCC1, and NKCC2 through a yeast two-hybrid screen and described the canonical RFXV motif in the cotransporters ^4,9^. Later on, SPAK and OSR1 were shown to be substrates of WNKs, and RFXV motifs within WNKs were described to mediate interactions with these downstream kinases ^2,3,10^.

Structural characterization showed that the CCT domain of OSR1 adopts a compact fold with an antiparallel β-sheet composed of four β-strands and two α-helices packed against it, helping stabilize the overall fold and contributing to the formation of the ligand-binding surface ^11^. Together, the β-sheet, the flanking helices, and the connecting loops create a shallow peptide-binding groove that recognizes short linear motifs containing the canonical sequence RFXV/I present in binding partners. A recent work by Taylor et al. performed a careful assessment of the binding and affinity of the CCT domains of SPAK and OSR1 to a peptide array derived from the CCT-binding peptide of hWNK4 ^12^. They confirmed that the CCT domains of SPAK and OSR1 recognize short linear motifs centered on the canonical R-F-x-V/I sequence (but also the related R-x-F-x-V/I variant) and concluded that although the core residues are essential for binding, the surrounding residues can modulate interaction affinity and specificity.

WNK kinases have also been shown to contain globular domains within their C-terminal region that adopt a fold similar to that of the CCTs of SPAK and OSR1 ^13–15^. From insects to mammals, WNKs contain two of these domains, termed CCT1 and CCT2, which are composed of a three-stranded β-sheet and two α-helices ^13,15^. This CCT1 was originally proposed to function as an autoinhibitory domain ^16^; however, more recent evidence indicates a different role, as described below.

NRBP proteins and TSC22D proteins were recently identified as interactors of WNK kinases and modulators of the WNK signaling pathway ^17–19^. Interestingly, the interaction of NRBPs with TSC22D proteins, as well as the interaction of TSC22Ds with WNKs and SPAK, appears to involve CCT domains and RFXV-like motifs ^12,17–19^. Specifically, NRBP1 and NRBP2, which are pseudokinases evolutionarily related to WNKs, contain CCT domains in their C-terminus that mediate binding to the long isoforms of the TSC22D family ^17,18^. These long TSC22D isoforms are proteins containing a long N-terminal intrinsically disordered region (IDR) and a short, folded domain at their C-terminus known as TSC22 domain. The TSC22 domain is responsible for protein homo- and heterodimerization ^18^. Within their N-terminal IDR, long TSC22D isoforms present a short, highly conserved region that contains two characterized CCT binding motifs, one of them containing the canonical RFRV sequence (termed RϕA by Amnekar et al. ^18^), which was shown to mediate binding to the CCT1 of WNKs, and another one displaying some divergence from the canonical sequence, being RWTC (termed RϕB). This latter motif was shown to mediate binding with NRBP1 ^18^.

The function of the CCT domains of WNKs was obscure until we recently described that mutation of key residues within both CCT domains of WNK4 disrupts its interaction with TSC22D2 and prevents activation of WNKs by the NRBP1/TSC22D complex ^19^. Moreover, Amnekar et al. showed that mutation of the RϕA motif in TSC22Ds decreased its binding to the isolated CCT1 domain of WNK1, but not to the CCT2 domain of WNK1 ^18^. Binding to neither of these domains was affected by mutation of the RϕB motif (RWTC), suggesting that this motif specifically mediates binding to NRBP1, and raising the possibility that a third motif may exist in TSC22Ds that mediates binding to the CCT2 of WNKs.

In the present study, we used structural modeling to identify the motif in TSC22Ds that binds to the CCT2 of WNK kinases. Interestingly, this motif differs from the canonical RFXV sequence but preserves features that highlight the most relevant interactions that govern domain-motif binding. We show that known CCT domains can be classified into four group based on structural and sequence resemblance, and that each of these groups displays a preference for different types of motifs, establishing specificity in the way that pathway components interact.

Finally, inspired by the high structural resemblance among different types of CCTs, we wondered whether we could identify additional mammalian proteins harboring potential CCT domains based on structural homology. We used the structural comparison platforms Dali and Foldseek to screen the Alphafold3 database and identified that only a handful of proteins present domains displaying a CCT-like fold. Among these proteins, we identified Ferry3, which is a component of the FERRY complex, a Rab5 effector that links early endosomes to the transport of specific mRNAs ^20–22^. Interestingly, the isolated CCT-like domain of Ferry3 can interact with TSC22D2, and this interaction depends on the CCT-binding motifs present in TSC22D2, suggesting that this domain may indeed function as a CCT. Future research will be necessary to establish the physiological binding partner that interacts with Ferry3 through this mechanism.

## Results

### Identification of the motif that mediates TSC22D binding to the CCT2 domain of WNKs

Structural and sequence comparison of different CCT domains revealed that these can be classified into four different groups (Supplementary Fig. 1): 1) a group containing the CCTs of SPAK and OSR1; 2) a group containing the CCTs of NRBP1 and NRBP2; 3) a group containing the CCT1 domains of WNKs; 4) and a group containing the CCT2 domains of WNKs. As mentioned, for each of these groups, binding motifs have been characterized, except for the CCT2 domain of WNKs, which has been shown to mediate interaction with TSC22D proteins ^19^.

Inspired by Amnekar’s et al work, in which it was shown that mutation of the RFxV and RWTC motifs in TSC22Ds (named RϕA and RϕB) did not affect binding to the CCT2 domain of WNK1 ^18^, we used AlphaFold3 to model the interaction between human WNK4 (Q96J92-1) and human TSC22D2 (O75157-1) to identify the binding motif responsible for this interaction. Interestingly, as suggested by experimental data, modeling did not predict binding of the CCT2 domain of WNK4 to the RFxV and RWTC motifs in TSC22D2, but predicted binding to a region located at the N-terminus of TSC22D2 (Fig. 1A). Remarkably, the identified binding region is highly conserved across species and is present in all TSC22D paralogues (Fig. 1B), suggesting functional relevance. The sequence logo for this newly identified motif highlights a conserved KxxxxFxITSV sequence (Fig. 1B), which in most vertebrates is present as KKKSxFQITSV. Notably, this binding region differs significantly from the canonical RFXV motif, but as explained below, it preserves certain key features.

**Figure 1.**
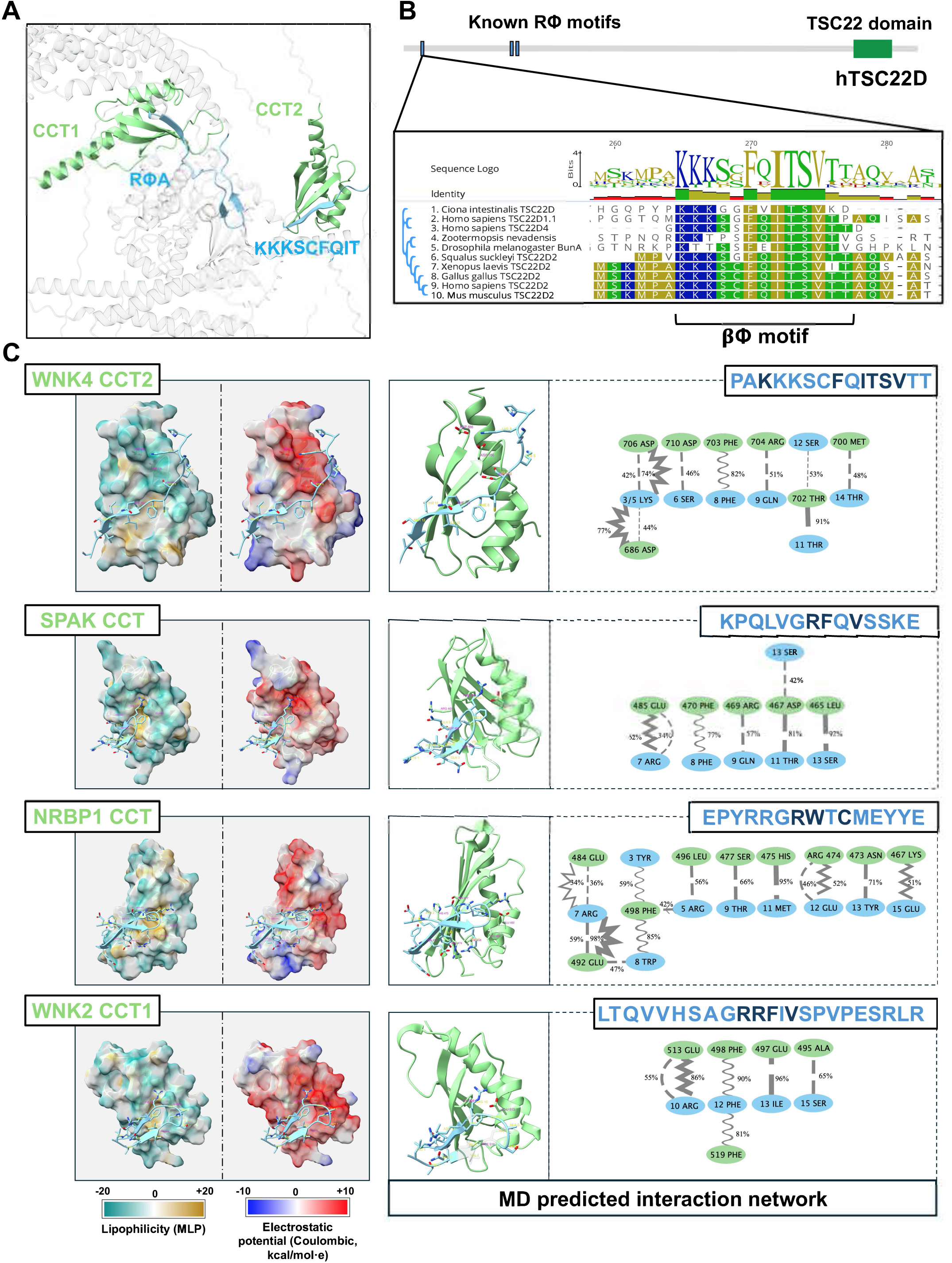
CCT2 domains of WNKs bind to a motif in TSC22D proteins that significantly differs from the canonical RFxV/I motifs bound by OSR1/SPAK CCT domains. **(A)** AF3-predicted dimer of hWNK4 and hTSC22D2. The sequences for human WNK4 (Q96J92-1) and human TSC22D2 (O75157-1) were used to model the interaction among these proteins in AF3. The highlighted interface regions had a pLDDT score >70%. **(B)** Sequence alignment of hTSC22D2 orthologues and paralogues. The sequence logo of the motif is depicted above and was generated using Geneious Prime 2026.0.2. **(C)** Representative structures at equilibrium from the molecular dynamics are presented for each of the modeled CCT domains. A CCT domain representative of each of the four identified CCT groups was selected for this analysis. The sequence of the binding peptides used for the simulations is shown in the top right corner of each panel for each CCT. For each CCT, calculated hydrophobic and electrostatic surfaces are shown to the left. All CCTs contain an acidic region in proximity to the positively charged residues of the bound peptide. To the right, the structures in ribbon display are shown and the residues involved in interaction with the CCT-binding peptides are highlighted. Also shown to the right are the diagrams depicting the most frequent interactions that were observed during the molecular dynamics. Numbering for the domain’s residues correspond to that of the entire protein, while numbering for the bound peptide only considers the position of the residue within the peptide. Dashed lines represent hydrogen bonds, wavy lines represent π-stack interactions, zigzag lines represent electrostatic interactions. Line thickness is proportional to the frequency in which each interaction was observed, and average frequency values are indicated. The characteristic fold of this type of domains can be appreciated: a β-sheet (composed of 4 β-strands in CCTs of SPAK/OSR1 and three β-stands in CCTs of WNKs and NRBP1) and two α-helices packed against the β-sheet. Between these structures the peptide binding groove is formed in which a central hydrophobic pocket accommodates the aromatic residue of the binding peptide, while electrostatic interactions help position the upstream positive residues and orient the peptide for stable binding.

Analysis of the interaction networks derived from molecular dynamics simulations of CCT domains in complex with bound peptides revealed that, in all cases, binding depends on two main anchoring modes: electrostatic anchoring and aromatic recognition (Fig. 1C, Supplementary Fig. S2-6, Supplementary table 1). When considering the most prevalent interactions (>20% occupancy), each complex showed that the peptide establishes at least one salt bridge and one hydrogen bond involving a basic residue within the peptide with an acidic residue in the CCT domain, consistent with an electrostatic anchoring mechanism. For the WNK4-CCT2 bound to the PAKKKSCFQITSVTT peptide, we observed that particularly Lys3 and Lys5, act as electrostatic anchors by forming ionic interactions and hydrogen bonds with the acidic residues Asp706 and Asp686 of the domain.

Regarding the second anchoring mode, for all CCTs, aromatic recognition was mediated by the peptide’s aromatic residue, which forms a π-stacking interaction with at least one aromatic residue of the CCT domain. In the case of the CCT2 of hWNK4, this residue corresponds to Phe703. Hydrophobic surface calculations of CCT-βφ peptide complex models revealed that, in all cases, the peptide’s aromatic residue lies within a hydrophobic pocket of the CCT. This pocket contains aliphatic residues and at least one aromatic residue capable of engaging in π-stacking interactions. The critical contribution of interactions involving the peptide basic residue (R) or aromatic residue (F or W) has been demonstrated previously in studies in which substitution of residues involved in these interactions destabilized the peptide–domain complex and abolished binding ^6,11,12,18,23,24^. Taken together, these results suggest that WNK4 CCT2 recognizes a motif with physicochemical features compatible with those of other CCT-binding peptides, although its sequence differs from those reported previously ^4,12^.

Sequence alignment of CCT domains revealed that many residues implicated in key interactions, based on molecular dynamics simulations and previous studies, are highly conserved across CCTs. However, notable exceptions exist. Although CCT2 domains retain residues that in other CCTs are involved in electrostatic anchoring (Supplementary Fig. 7, green arrows), it contains two additional downstream aspartate residues (Asp686 and Asp706, Supplementary Fig. 7, red arrows) that mediate this interaction. These residues are not fully conserved in other CCTs, with the second aspartate (Asp706) conserved only in CCT1. This feature may explain CCT2’s ability to accommodate peptides in which lysine residues are positioned farther upstream of the motif’s central phenylalanine.

Also notorious is the lack of conservation of the phenylalanine residue involved in π-stacking interactions, which is present in WNKs CCT1, CCT2, and SPAK (e.g. Phe452 in hOSR1), but absent in NRBPs (Supplementary Fig. 7, first blue arrow). In NRBP, this phenylalanine is instead located further downstream, at a position where a phenylalanine or leucine is found in other CCTs that hass also been implicated in peptide interaction (Supplementary Fig. 7, second blue arrow, e.g., Leu473 in hOSR1, Leu502 in mSPAK) ^11,23^.

Previously, Amnekar et al. proposed naming the CCT-binding motifs as Rϕ motifs (ϕ for hydrophobic) with basis in the observation that NRBP1 could accommodate a tryptophan-containing peptide within the CCT’s hydrophobic groove. However, in light of the recent observation that even a more divergent binding motif is present in peptides that bind the CCT2 of WNKs, we propose to call them βϕ motifs (β for positively charged and ϕ for hydrophobic), to highlight the residues involved in the electrostatic and aromatic anchoring, features that appear to be present in all CCT-binding motifs identified so far.

### Each type of CCT domain displays distinct binding preference for different βϕ motif types

We used Alphafold3 to model the interaction of distinct types of CCTs bound to five different known βϕ-motif containing peptides. Fig. 2A shows the iPTM values for each complex. These values reflect the accuracy of the interface predictions. From this analysis, we observe a tendency toward higher ipTM values in some cases, suggesting that distinct types of CCTs prefer different βφ motif types.

**Figure 2.**
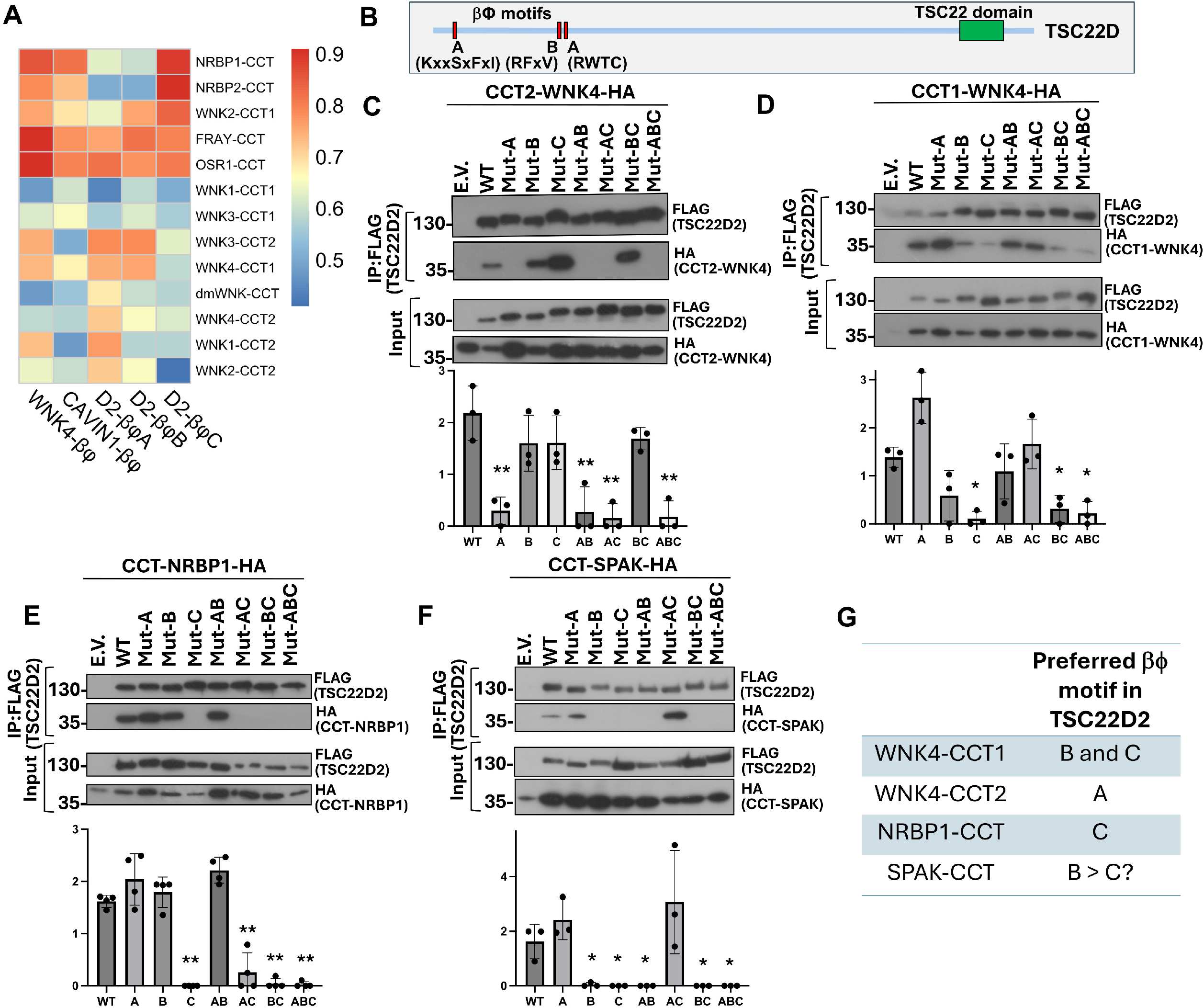
Preferred binding motifs for different types of CCTs. **(A)** Heat map of iPTM values obtained for each AF3 model of the indicated CCT-βϕ complex. **(B)** Diagram depicting the position of the three βϕ motifs present in long TSC22D isoforms. **(C-F)** Assessment of the effect of mutations in the βϕ motifs of TSC22D2 on the interaction with the CCT2 domain of WNK4 (C), the CCT1 domain of WNK4 (D), the CCT domain of NRBP1 (E), or the CCT domain of SPAK (F). At least three independent experiments were performed. Results of quantification of the bound co-immunoprecipitated CCT are shown in the graphs under each representative blot (mean ± SD). ANOVA followed by Tukey’s post hoc tests were performed to identify statistically significant differences. ^*^P < 0.05, ^**^P<0.001. **(G)** According to the results presented in panels C-F, the preferred βϕ motif in TSC22D proteins bound by the different types of CCTs are enlisted.

In order to validate the newly identified βϕ motif as a CCT2 binding motif, as well as to experimentally compare the binding preference of the different types of CCT domains, we performed co-immunoprecipitation experiments in which selected isolated CCTs (representing each of the four groups of CCTs, Supplementary Fig. 1) were co-expressed with wild-type TSC22D2 or with TSC22D mutant constructs in which the βϕ motifs were individually or collectively mutated. We decided on this strategy because TSC22Ds contain the canonical CCT-binding motif, RFxV, as well as the two alternative CCT-binding motifs described so far (KKKSxFQITSV and RWTC) (Fig. 2B).

As expected, mutation of the βϕA motif of TSC22D2 prevented binding to the isolated CCT2 domain of WNK4, whereas individual mutation of the βϕB or βϕC motifs did not prevent binding (Fig. 2C). In addition, the presence of the βϕA motif in the absence of the βϕB and βϕC motifs (Mut-BC) was sufficient to observe interaction with the CCT2 domain of WNK4. This confirms that the βϕA motif is a CCT-binding motif that selectively binds CCT2 motifs present in WNKs.

For the isolated CCT1 of WNK4, we observed that individual mutation βϕA of TSC22D2 did not affect the interaction with this CCT, whereas individual mutation of the βϕB or βϕC motifs dramatically decreased the interaction with this CCT (Fig. 2D). Consistently, having only the βϕB (Mut-AC) or the βϕC (Mut-AB) motifs was sufficient to observe interaction, and the construction having only the βϕA motif was unable to interact. This slightly contrasts with what was reported by Amnekar et. al., who observed that mutation of the βϕB, but not the βϕC of TSC22D2 or TSC22D4, reduced the interaction with the isolated CCT1 of WNK1 ^18^.

For the isolated CCT of NRBP1, all TSC22D2 constructs lacking the βϕC motif (Mut-C, Mut-AC, Mut-BC, and Mut-ABC) showed dramatically reduced interaction, whereas having only the βϕC motif was sufficient to maintain interaction (Mut-AB) (Fig. 2E). This is consistent with previous data by Amnekar et al. showing that the NRBP1 interacts with TSC22Ds through the βϕC motif ^18^.

Finally, for the CCT of SPAK, individual mutation of motifs βϕB or βϕC reduced interaction, whereas mutation of the βϕA motif did not affect it (Fig. 2F). The presence of only the βϕB motif (Mut-AC) was sufficient to observe interaction. This is consistent with the knowledge that SPAK and OSR1 CCTs preferentially bind peptides containing the RFxV consensus sequence ^4,5,12^. However, no interaction was observed with the construct harboring only the βϕC motif (Mut-AB). Supporting this observation, Taylor et al. reported that substitution of canonical phenylalanine with tryptophan in the RFxV motif is poorly tolerated by the SPAK/OSR1 CCT domain. Therefore, the reduced interaction observed upon mutation of the βϕC motif may reflect indirect effects on binding rather than a primary contribution of this motif.

In summary, these results suggest that among the different βϕ motifs present in TSC22D proteins, the βϕA motif mediates interaction with the CCT2 of WNKs, the βϕB motif can mediate the interaction with the CCT1 of WNKs as well as the CCT of SPAK, and the βϕC motif can mediate interaction with NRBP1, as well as with the CCT1 of WNK1 (Fig. 2G). Thus, TSC22D proteins serve as scaffolds that bring together components of the WNK pathway through divergent βϕ motifs that mediate interactions with various CCT domains.

### Analysis of the interaction of full-length proteins with TSC22D mutants

We next assessed the interactions between the different TSC22D mutant constructs and the full-length WNK4, NRBP1, or SPAK proteins. In accordance with the binding preference observed for the isolated CCTs of WNK4, we observed that TSC22D2 only lost the interaction with full length WNK4 when the βϕA and βϕB were mutated simultaneously, supporting the concept that these motifs interact with the CCT2 and CCT1 domains of WNKs, respectively and that presence of at least one motif is sufficient to maintain the interaction (Fig. 3A, D).

**Figure 3.**
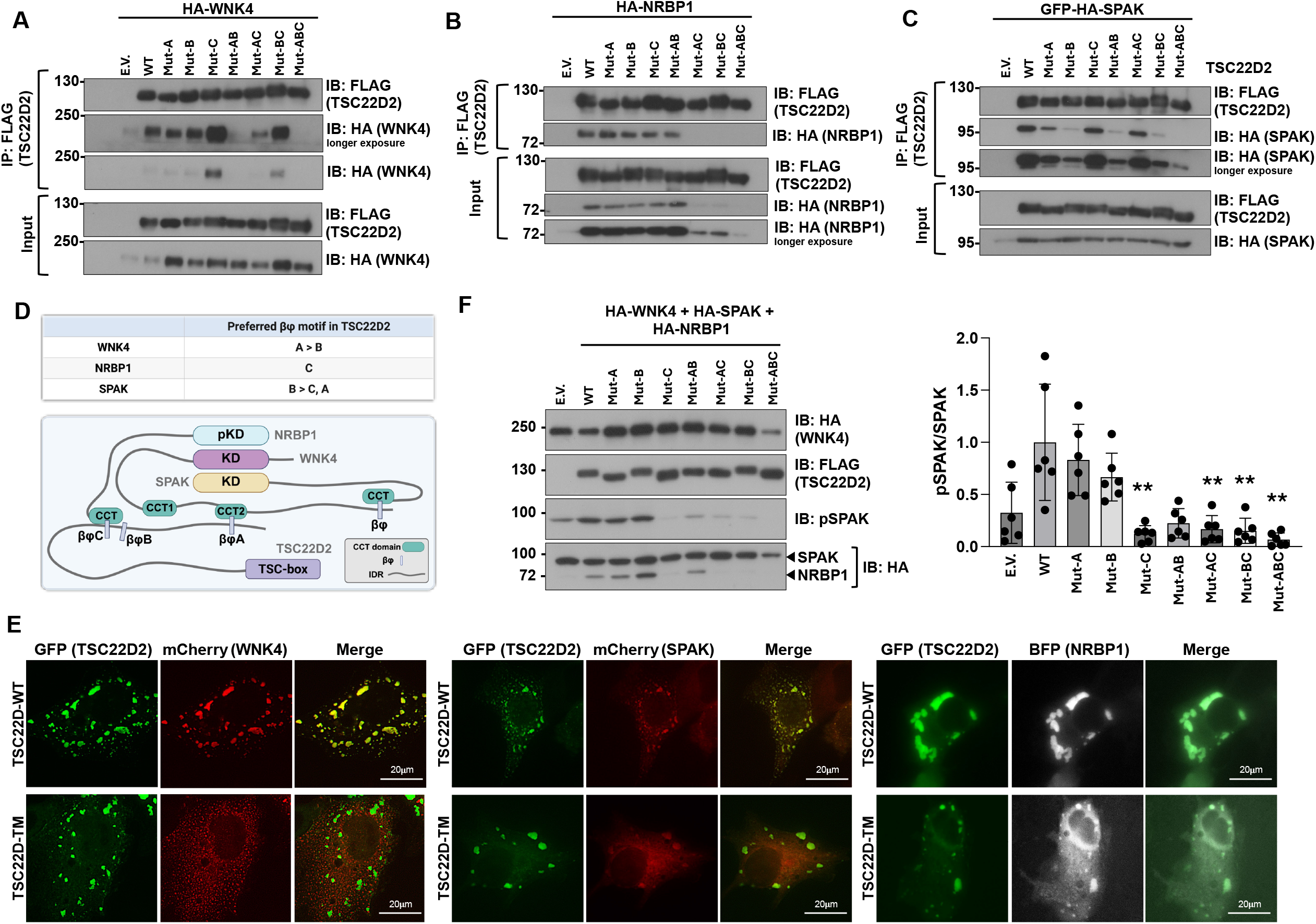
Effect of mutations of the βϕ motifs in TSC22D2 on its binding to WNK4, NRBP1 or SPAK. **(A-C)** Assessment of the effect of mutations in the βϕ motifs of TSC22D2 on its interaction with WNK4 (A), NRBP1 (B), or SPAK (C). **(D)** Summary of the binding preference for each of the analyzed proteins according to the results presented in A-C. **(E)** Assessment of colocalization of TSC22D2 and TSC22D-TM with WNK4, SPAK and NRBP1. COS7 cells were transiently transfected with WNK4-mCherry, SPAKmCherry, or NRBP1-BFP, together with TSC22D2GFP or TSC22D-TM-GFP. The degree of colocalization was quantified in Fiji and is presented in Supplementary Fig. 8. **(F)** Effect of disruption of the βϕ motifs on the ability of TSC22D2 to upregulate WNK-SPAK signaling. Six independent experiments were performed. Results of quantification of pSPAK signal intensity are shown in the graph to the left (mean ± SD). ANOVA followed by Tukey’s post hoc tests was performed to identify statistically significant differences. **P < 0.001.

For NRBP1, we observed that the TSC22D2 construct containing only the βϕC motif (Mut-AB) was able to interact with NRBP1, whereas the constructs containing only the βϕA (Mut-BC) or βϕB (Mut-AC) did not interact (Fig. 3B, D). Interestingly, as shown previously ^19^, interaction of TSC22D2 with NRBP1 was necessary to stabilize the NRBP1 protein, as higher NRBP1 abundance was observed in the presence of the TSC22D constructs with which interaction was observed. Surprisingly, in contrast to what was observed with the isolated CCT of NRBP1, interaction of full-length NRBP1 was observed with the TSC22D2 βϕC mutant (Mut-C). Although we cannot currently explain this observation, the explanation is necessarily related to the additional protein regions present in the full-length protein (e.g., the pseudokinase domain).

With respect to SPAK, we observed that the TSC22D2 βϕ mutation that had the greatest effect on SPAK binding was the βϕB mutation (Mut-B) (Fig. 3C, D). Mutation of the βϕA motif only slightly reduced interaction, whereas mutation of the βϕC motif had no effect. In addition, the TSC22D construct containing only the βϕB motif (Mut-AC) showed similar binding to SPAK as the wild-type protein, and the constructs containing only the βϕA (Mut-BC) or βϕC (Mut-AB) motifs showed reduced binding. These results are consistent with the preference of the CCT of SPAK for RFxV motifs reported previously ^4,9,12,24^.

We also assessed the effect of the absence of βϕ motifs in TSC22D2 on its colocalization with WNK4, SPAK, and NRBP1. WNK kinases have been shown to form biomolecular condensates via liquid-liquid phase separation in response to the molecular crowding that occurs when the cells are exposed to a hypertonic stimulus ^25^. Within these condensates, other components of the signaling pathway are also present, like SPAK/OSR1, TSC22D, and NRBP proteins ^19,25,26^. In our overexpression system, we have previously reported that WNK condensates can be observed under isotonic conditions ^19^. Thus, as expected and previously observed, the wild-type TSC22D2-GFP colocalized with WNK4-mCherry, SPAK-mCherry, and NRBP1-BFP in condensates (Fig. 3E and Supplementary Fig. 8). In contrast, WNK4 and TSC22D2-TM were clearly distributed to distinct subcellular compartments, i.e., distinct WNK4-positive and TSC22D2-positive condensates were formed. Similarly, reduced colocalization of SPAK with the TSC22D2-TM was observed. Interestingly, however, despite the decreased interaction observed in the immunoprecipitation experiments, colocalization of the TSC22D2-TM with NRBP1 was still observed.

In cells cotransfected with WT-TSC22D2-GFP and WNK4-mCherry, stimulation with hypertonic stress (Supplementary Fig. 9) increased the number of condensates in which WNK4 and WT TSC22D2 perfectly colocalized. In contrast, in cells co-expressing WNK4-mCherry and TSC22D2-TM-GFP, an increase in WNK4-positive condensates was observed upon hypertonic stimulation, but no increase in the formation of TSC22D2 condensates was observed. This observation suggests that, although both WNK kinases and TSC22D proteins possess IDRs and have been shown to promote condensation ^19^, WNK kinases appear to be the major drivers of condensate formation in response to hypertonicity.

Finally, we assessed the effect of mutations in the βϕ motifs on TSC22D2’s ability to upregulate the activity of the WNK-SPAK pathway. As previously reported ^19^, expression of wild-type TSC22D2 in the context of WNK4, SPAK, and NRBP1 overexpression increased the levels of pSPAK (Fig. 3F). pSPAK levels observed in the presence of the TSC22D2 constructs harboring individual mutations in the βϕA (Mut-A) or βϕB (Mut-B) motifs were similar to that observed with wild-type TSC22D2. However, in the presence of the βϕC mutant (Mut-C), a significantly lower level of pSPAK was observed. Additionally, in the presence of all double mutants, reduced pSPAK levels were observed. Thus, the absence of the βϕC, which is mainly involved in interaction with NRBP1, significantly affects the ability of TSC22D proteins to upregulate WNK signaling, and the absence of βϕA or βϕB motifs only has a significant effect when mutated in conjunction with other βϕ motifs.

### Structural homology-based search for other CCT-containing proteins identifies a handful of proteins that harbor globular domains with a CCT-like fold

We next wondered whether other proteins in the human proteome contain domains with a CCT-like fold that could mediate interactions. Finding such domains could potentially help us identify other components of the WNK pathway or proteins with unrelated functions that have adopted this interaction mechanism. To search for these domains in the human proteome, we used the structural comparison platforms Dali ^27,28^ and Foldseek ^29^ and screened the Alphafold3 database (Fig. 4A and B).

**Figure 4.**
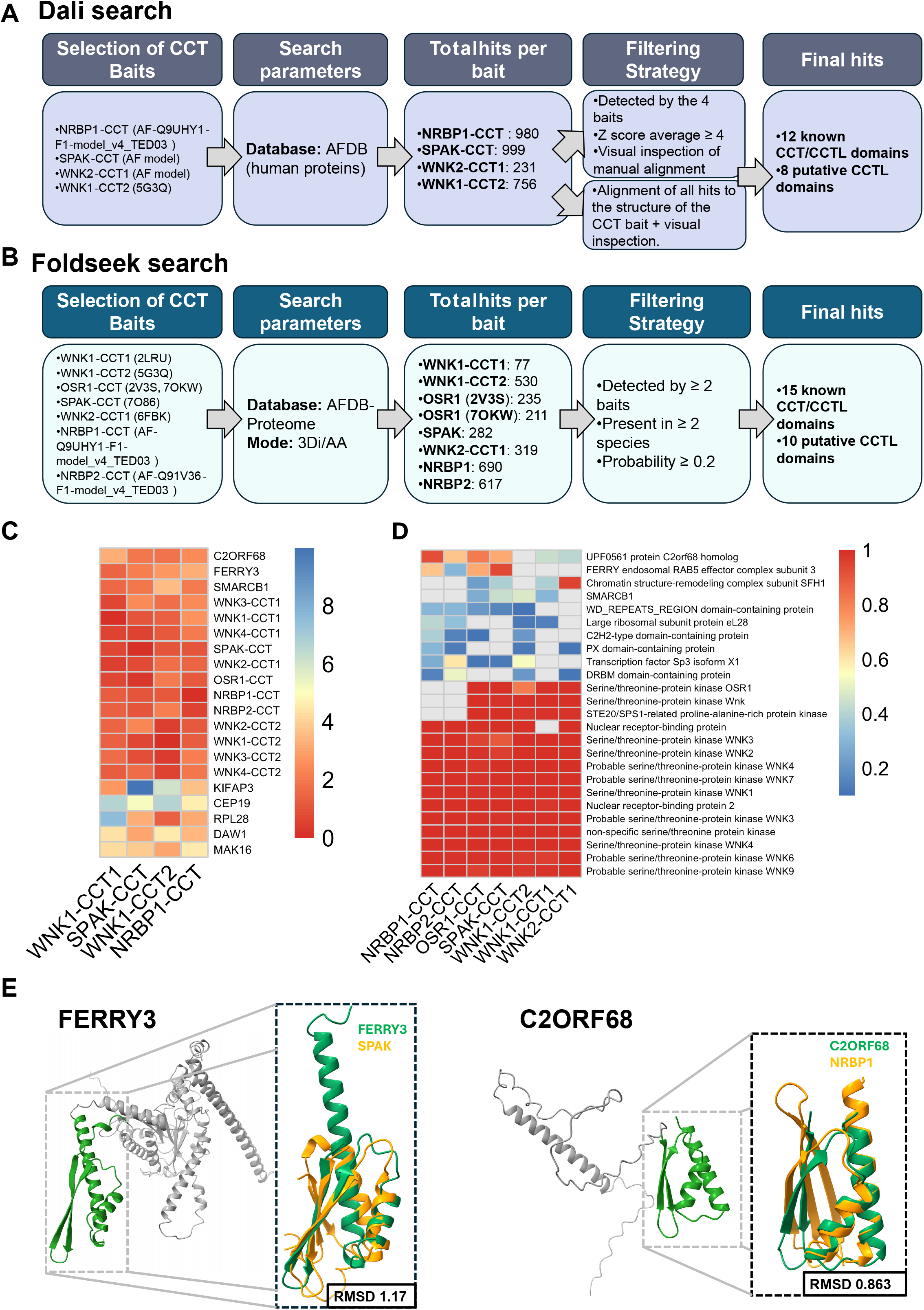
Structure-guided search of CCT-like domains in the human proteome. **(A)** DALI search. Workflow of the structure-based search using DALI to identify CCT/CCTL domains. This approach yielded a set of known CCT domains and additional putative candidates. **(B)** Foldseek search. Complementary structure-based search performed using Foldseek against the AlphaFold proteome database. This strategy confirmed the top candidates found with DALI (FERRY3 and C2ORF68) and uncovered additional putative domains. **(C)** Heatmap of Z scores indicating the resemblance with each of the CCT baits for the top hits obtained from the DALI search. Known CCT-containing proteins (e.g., WNKs, SPAK/OSR1, NRBP1/2) show strong and consistent detection across baits, whereas other proteins display more variable or selective detection patterns, supporting their classification as putative CCT-like-containing proteins. **(D)** Heatmap summarizing structural similarity scores (Foldseek probability) between identified hits and canonical CCT domains. Known CCT-containing proteins cluster with high similarity, while putative candidates, particularly FERRY and C2ORF68 display intermediate but significant structural resemblance. **(E)** Alphafold3 model structures for FERRY3 and C2ORF68. The region of the protein which was identified to have a CCT-like fold is highlighted in green. Insets show structural superposition of predicted domains from FERRY3 and C2ORF68 with the indicated canonical CCT domains. The low RMSD values highlight the strong structural resemblance despite sequence divergence.

For the Dali search, four known CCT domains were used as queries (Fig. 4A). On average, each CCT query yielded 517 hits. These hit lists were merged and curated to retain only proteins detected by all four baits and with Dali Z-scores ≥ 4 (Supplementary table 2). In parallel, and to confirm that the filtering strategy did not lose potential candidates, manual alignment of all hits to the corresponding CCT bait was performed, and potential candidates were selected by visual inspection of the alignment. As shown in Fig. 4C this strategy was sensitive enough to detect all 12 known CCT domains that have been described in the human proteome (Supplementary Fig. 1). These served as positive controls. In addition, 8 putative CCT-like domains were found in the proteins encoded by C2ORF68 (Q2NKX9), FERRY3 (Q9NQ89), SMARCB1 (Q12824), KIFAP3 (Q92845), CEP19 (Q96LK0), RPL28 (P46779), DAW1 (Q8N136) and MAK16 (P10962).

For the Foldseek search, eight known CCT domains were used as queries (Fig. 4B). On average, each CCT query yielded 370 hits (Supplementary table 3). The hit lists were merged and curated by retaining only proteins detected by two or more baits, that were present in at least two species and that had an average probability score for all detected orthologues ≥ 0.2. Given that the database searched here was the AFDB-proteome, which also include non-human proteins, in this case we found more than 12 proteins containing known CCT domains (Fig. 4D). In addition, 10 putative CCT-like domains were found in the proteins encoded by C2ORF68 (Q2NKX9), FERRY3 (Q9NQ89), SFH1 (A0A1D8PEB6), SMARCB1 (Q12824), RPL28 (P46779), WDSUB1 (Q8N9V3), PXDC1 (Q5TGL8), SP3 (Q02447), attf-3 (G4S460) and Tb03.28C22.80 (Q57WD9).

Interestingly, the two putative CCT-like domains that showed higher similarity scores to known CCTs under both search strategies were those present in the proteins C2ORF68 and FERRY3. The structure of these proteins has not been experimentally determined. However, inspection of the Alphafold3 model structures for these proteins revealed that the regions of the proteins that were identified to have a CCT-like fold correspond to modular globular domains (AF-Q9NQ89-F1-model_v4_TED01 for FERRY3 and AF-Q2NKX9-F1-model_v4_TED02 for C2ORF68) connected to the rest of the protein by disordered regions (Fig. 3E). This contrasted with the analysis of other hits, where the putative CCT-like domain appears to be part of a larger globular domain that probably has a different function (e.g. SMARCB1).

### Ferry3 contains a CCT-like domain that can mediate interactions with βϕ -like motifs

Analysis of structural conservation of the putative CCT domains of FERRY3 and C2ORF68 in distant orthologues’ models revealed a high degree of conservation, suggesting that this fold is relevant to the function of these domains (Fig. 5A, C). Conservation was also observed at the primary sequence level (Fig. 5A, C)

**Figure 5.**
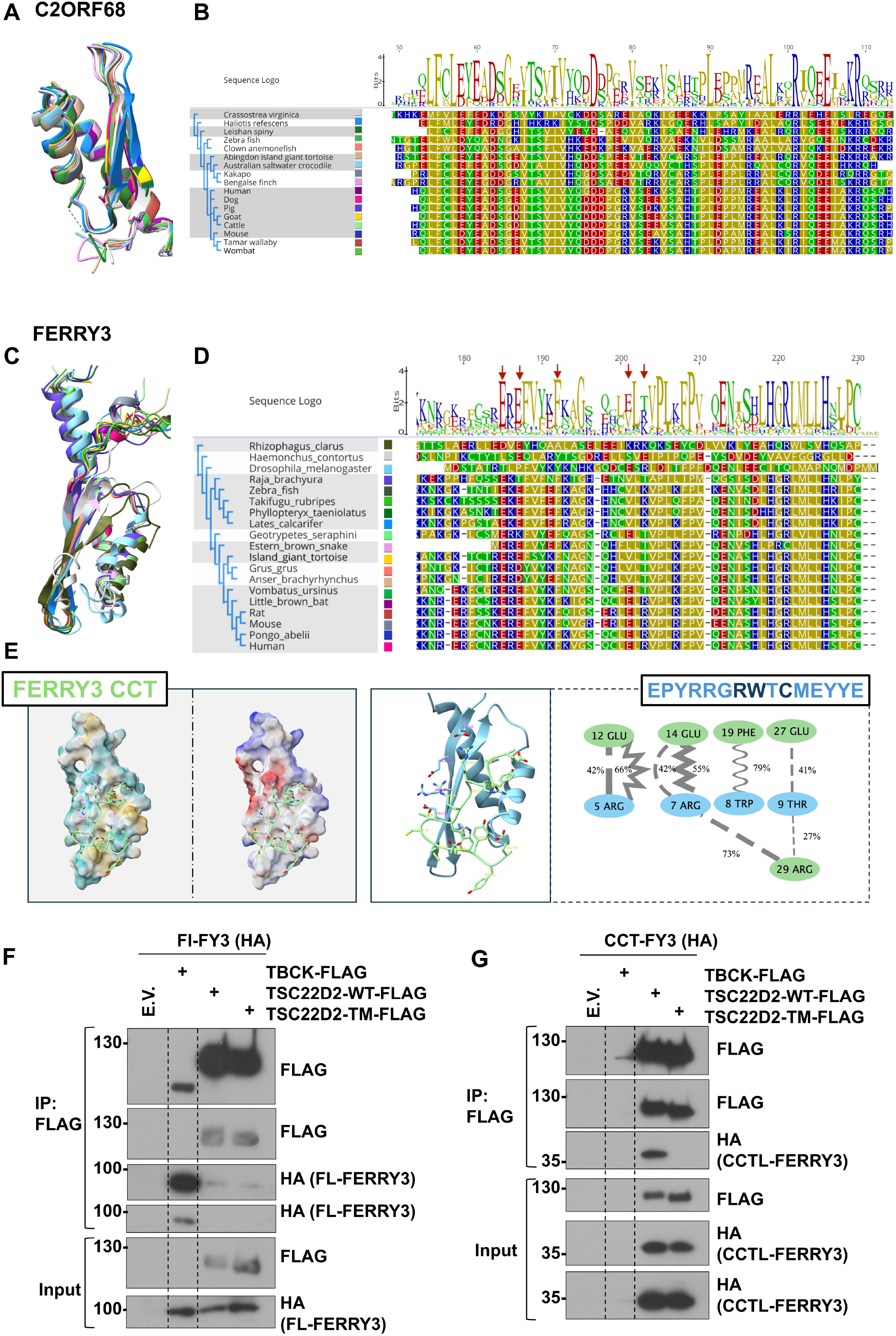
Structural conservation and functional assessment of the putative CCT domain in FERRY3. **(A, C)** Structural conservation of the putative CCT domains of C2ORF68 and FERRY across distant orthologues, showing preservation of the overall fold. **(B, D)** Corresponding sequence conservation is also observed at the primary sequence level. **(E)** Molecular dynamics simulations of the FERRY3 putative CCT domain in complex with a βϕC-containing peptide. The peptide binds to a groove formed by the second β-strand and the adjacent α-helix, consistent with canonical CCT–βϕ interactions. Electrostatic anchoring (Arg5/Arg6 with Glu12/Glu14) and aromatic stacking (Trp8 with Phe19) are observed with high frequency. **(F, G)** Co-immunoprecipitation analysis of FERRY3 interactions. HA-tagged full-length FERRY3 (F) or the isolated putative CCT of FERRY3 (G) were overexpressed in HEK293 cells together with Flag-tagged WT TSC22D2, TSC22D-TM, or TBCK. Flag IP was performed and co-immunoprecipitations assessed by immunoblot. Full-length FERRY3 does not interact with TSC22D2, whereas the isolated putative CCT domain interacts with TSC22D2 but not with the TSC22D2-TM mutant lacking bf motifs. Three independent experiments were performed with similar results.

As a preliminary analysis of their potential to function as CCTs, their interaction with the three βϕ motifs of TSC22D2 (which represent the three types of βϕ motifs known) was modeled with Alphafold3 (Supplementary Fig. 10). Acceptable iPTM values were obtained only for the FERRY3-βϕC complex and the C2ORF68-βϕB complex. However, visual inspection of the latter complex revealed that the βϕ motif peptide bound to a cavity of the C2ORF68 putative CCT that does not correspond to the hydrophobic groove in which these motifs normally bind.

Guided by these results, we performed molecular dynamics simulations of the FERRY3 putative CCT domain in complex with the βϕC motif-containing peptide (Fig. E and Supplementary Fig.11). Interestingly, in these simulations, the βϕC peptide bound to a groove formed by the second beta stand and the subsequent alpha helix, as it occurs in known CCT-βϕ complexes. In addition, the two characteristic anchoring modes that are known to direct binding of known CCT-βϕ complexes (electrostatic anchoring and aromatic recognition) were also observed to occur with a frequency >50%. In particular, Arg5 and Arg6 of the peptide established salt bridges and hydrogen bonds with Glu12 and Glu14 of the putative CCT, respectively, and Trp8 established π-stack interactions with Phe 19 in the FERRY-CCT. Notably, these residues are highly conserved (Fig. 5B).

Finally, FERRY3 was cloned, and its potential interaction with TSC22D2 was assessed by coimmunoprecipitation. The putative CCT domain in FERRY may not necessarily mediate interaction with a component of the WNK pathway and may instead participate in an interaction with a binding partner that is relevant to the role of this protein in the cell. However, given that this binding partner is currently unknown, we chose TSC22D2 to assess interaction because it contains the three known types of βϕ motifs, including the βϕC motif, which can potentially mediate interaction according to in silico simulations. We reasoned that at least a low-affinity interaction could be observed if one of these motifs is similar to the putative physiological binding motif. No interaction between full-length FERRY3 and TSC22D2 was observed in coimmunoprecipitation experiments performed with overexpressed proteins in HEK293 cells (Fig. 5F). Interaction with the known FERRY3 binding partner, TBCK, a known interacting partner ^20,21^, served as a positive control.

Given that FERRY3 has been shown to function as part of a complex that directs mRNA transport and that is associated with early endosomes ^20,21^, we reasoned that perhaps the lack of detectable interaction could be due to a lack of colocalization. Thus, we decided to assess the interaction between TSC22D2 and the isolated putative CCT domain of FERRY3 (Fig. 5G). Interestingly, the isolated putative CCT of FERRY was unable to interact with TBCK, suggesting that this interaction is mediated by another region of the protein, but was able to interact with TSC22D2. Importantly, interaction was not observed with the TSC22D2-TM construct, which lacks the three βϕ motifs, suggesting that interaction with the wild-type protein was mediated by at least one of these motifs. Future studies will aim to identifying the binding partners of FERRY3 that could interact through this putative CCT domain.

## Discussion

CCT domains have traditionally been viewed as modules that recognize canonical RFxV motifs ^4,5,12^; however, our findings indicate that this model captures only a subset of CCT-mediated interactions. We show that CCT domains can be classified into distinct structural groups that exhibit selective binding to divergent βφ motifs, thereby establishing a specificity framework based on CCT–motif pairing within the WNK signaling pathway. This classification helps reconcile previous observations that could not be fully explained by canonical RFxV-based recognition alone, including the selective binding of NRBP proteins to RWTC-like motifs ^18^ and the lack of interaction of WNK CCT2 domains with canonical motifs ^18,30^. Interestingly, an analogous scenario has been described for WW domains, which have been classified according to structural features that define their binding specificity ^31^.

Our work suggests that despite substantial sequence divergence among the newly identified βφ motifs, binding converges on shared physicochemical principles. Across all complexes analyzed, interactions are consistently supported by electrostatic anchoring of basic residues and aromatic recognition within a hydrophobic pocket. Structural differences among CCT domains appear to reposition these interaction determinants, thereby enabling selective recognition of distinct motif architectures. In this context, the βφ nomenclature proposed here expands the definition of CCT-binding motifs by emphasizing conserved physicochemical features rather than strict sequence conservation.

These findings have important implications for how signaling specificity is achieved within the WNK pathway. TSC22D proteins contain multiple βφ motifs that engage different classes of CCT domains, supporting a model in which they function as modular scaffolds that integrate pathway components. Distinct combinations of interactions can be established depending on motif availability and CCT domain identity. In addition, differential specificity among CCT–βφ motif complexes may underlie the observation that certain interactions among pathway components are stimulated by hypertonicity whereas others are not ^17,18^. It remains to be determined whether the formation of a structurally organized complex, in which specific interactions among components occur, is required for enzymatic activity, or whether these interactions are more flexible and primarily serve to bring components into proximity.

It is noteworthy that plant WNK kinases contain a single CCT-like domain, whose functional role remains undefined. Structurally, this domain most closely resembles the CCT domains of metazoan NRBP proteins (Supplementary Fig. XX). Importantly, key residues implicated in β–φ motif recognition in other CCT domains are conserved, suggesting that plant WNK CCTs may retain the capacity to engage related peptide motifs (Supplementary Fig. 7). However, canonical components of the metazoan WNK signaling network—including SPAK/OSR1 kinases, as well as NRBP and TSC22 family proteins—are absent in plants, indicating that any CCT-mediated interaction network in this lineage must involve distinct partners. Together, these observations support the idea that plant WNK CCT domains function as peptide-recognition modules, although their binding partners and motif specificity remain to be defined.

In addition to defining interaction specificity within the WNK pathway, our study suggests that CCT-mediated recognition may extend beyond this signaling network, although likely limited to a small number of proteins. The identification of a CCT-like domain in FERRY3 indicates that this structural module is present in at least one additional protein not previously linked to the WNK pathway. FERRY3 is a component of the FERRY complex, a Rab5 effector that connects early endosomes with the transport of specific mRNAs and has been reported to interact with proteins such as TBCK ^20–22^. Our data suggest that the CCT-like domain in FERRY3 is not responsible for the interaction with TBCK, and future studies will be required to define its functional role. Interestingly, although the structure of several components of the FERRY complex has been determined by cryo-EM ^21^, FERRY3 remains one of the unresolved components, and thus its interaction landscape is not yet defined.

Finally, our data indicate that the isolated CCT-like domain of FERRY3 can interact with TSC22D2 in a motif-dependent manner, supporting its potential to act as a CCT module. In contrast, no interaction was detected with the full-length protein under the conditions tested. This discrepancy may reflect constraints imposed by subcellular localization or conformational context, as FERRY3 is associated with early endosomes, whereas TSC22D2 may not efficiently access this compartment in the experimental system. Thus, it is possible that the physiological binding partners of FERRY3 differ from those tested here and that TSC22D2 provides only a proxy for βφ-like motif recognition.

## Materials and Methods

### Structural dendrogram of CCT domains

Pairwise structural comparisons among the CCT domains of WNK, SPAK, OSR1, and NRBP proteins were performed using the DALI server ^27,28^. The resulting analysis organized the domains by relative three-dimensional proximity, identifying clusters of structurally related domains. The dendrogram was interpreted as a representation of structural similarity rather than a strict phylogenetic relationship.

### Molecular Modeling

Structural models of the various CCT domains were generated using AlphaFold3 ^32^ and refined with the Rosetta Relax protocol on the ROSIE server ^33,34^. Model geometry and stereochemical quality were evaluated using MolProbity ^35^. In parallel, complexes between CCT domains and βφ peptides were modeled with AlphaFold3 ^32^, refined using the PeptDock/FlexPepDock protocol on the ROSIE server ^33,36^, and globally relaxed with the Rosetta Relax protocol ^33,34^. The final complex models were also assessed for geometry and stereochemical quality using MolProbity ^35^.

### Structural visualization and alignment

PDB structures were visualized in PyMOL version 3.1.6.1 and UCSF ChimeraX version 1.11. Structural alignments were performed in ChimeraX using MatchMaker for both visualization and manual inspection. Pairwise structural superpositions were conducted in PyMOL using the super command, and the resulting RMSD values were used to compare putative CCT domains. The super command was chosen for its ability to perform sequence-independent structural superposition, which is suitable for comparing proteins with limited sequence similarity or uncertain evolutionary relationships.

### Amino acid sequence alignment

Protein amino acid sequences were sourced from UniProt, Ensembl, and OrthoDB. Multiple sequence alignments were generated using the MUSCLE algorithm in MEGA version 12.1.2, then examined and visualized in Geneious Prime 2026.0.2.

### Molecular Dynamics

Molecular dynamics simulations were conducted for CCT–βφ peptide complexes. The crystallographic structures of the OSR1 CCT–GRFQVT complex (PDB: 2SV3) and the WNK2 CCT–LTQVVHSAGRRFIVSPVPESRLR complex (PDB: 6FBK) were analyzed, along with AlphaFold3-derived models of the SPAK CCT–KPQLVGRFQVTSSKE, WNK4 CCT2– PAKKKSCFQITSVTT, and NRBP CCT–EPYRRGRWTCMEYYE complexes. All-atom molecular dynamics simulations were performed using OpenMM 8.2 ^37^ on NVIDIA RTX 5080 GPUs, with the AMBER ff14SB force field for protein parameterization ^38^. Each system was solvated with a 1.2 nm padding at pH 7.4 using TIP3P water molecules ^39^ and neutralized with NaCl at 0.15 M. Following two-stage energy minimization, systems were heated from 0 to 300 K over 500 ps under the NVT ensemble and equilibrated for 2 ns under the NPT ensemble at 300 K and 1 bar using a Langevin integrator (friction coefficient 1 ps−1) and a Monte Carlo barostat, with positional restraints gradually released. Covalent bonds involving hydrogen atoms were constrained using OpenMM’s internal algorithms, and water molecules were treated as rigid. Long-range electrostatics were calculated using the particle mesh Ewald method ^40^ with a 10 Å cutoff. Hydrogen mass repartitioning ^41^ was applied by assigning a hydrogen mass of 4 amu, enabling a 4 fs integration timestep. Two independent 200 ns trajectories were generated for each system under the NPT ensemble, resulting in a total sampling time of 400 ns per system.

Molecular dynamics trajectories were analyzed using MDTraj ^42^. Global structural stability was assessed by calculating RMSD relative to the reference conformation over the entire simulation. Residue-level flexibility was quantified using Cα RMSF from self-aligned trajectories to eliminate global rotational and translational motion. Solvent-accessible surface area (SASA) was calculated per residue using the Shrake–Rupley algorithm in MDTraj. These metrics characterized global stability, local flexibility, and solvent exposure dynamics throughout the simulations. Custom Python scripts were developed with assistance from GitHub Copilot.

### Interaction network analysis of CCT–βφ complexes from molecular dynamics simulations

Each molecular dynamics simulation was performed in duplicate and extended to 200 ns. Only the 50–150 ns segment of each trajectory was used for residue–interaction network analysis, as all systems reached equilibrium before 50 ns according to trajectory stability analyses. This interval provided a homogeneous and computationally manageable sampling strategy and was considered the equilibrated phase for comparative analysis. For each replica, two non-overlapping temporal windows (50–100 ns and 100–150 ns) were defined, and each window was subdivided into three independent ensembles of 100 uniformly sampled conformations, resulting in six ensembles per replica. Residue–interaction networks were constructed for each ensemble using RING ^43^, with interactions assigned according to the geometric criteria implemented in the server: hydrogen bonds (donor–acceptor distance ≤ 3.9 Å, hydrogen–acceptor distance ≤ 2.5 Å), ionic interactions (charged group distance ≤ 4.0 Å), π–π stacking (ring-center distance ≤ 6.5 Å), π–cation interactions (cation–ring center distance ≤ 5.0 Å), π–hydrogen interactions (donor–ring center distance ≤ 4.3 Å), van der Waals contacts (van der Waals radii intersection fraction ≥ 0.01), disulfide bonds (S– S distance ≤ 2.5 Å), and metal-coordination interactions (metal–ligand distance ≤ 2.8 Å). These analyses produced six independent networks per replica. Networks were integrated into a replica-specific consensus network, with support across networks, mean interaction frequency, and mean interaction distance calculated for each edge. A mean interaction frequency threshold of 20% was applied to retain recurrent contacts while allowing detection of less frequent but reproducible interactions. Network analysis and visualization were performed in Cytoscape v3.10.4. The two consensus networks for each CCT–peptide complex were compared, and only interactions conserved between both replicas were retained for representation and downstream analysis.

### Fast-cloning and construct generation

All constructs were generated by Fast Cloning ^44^. Fast cloning was performed using the Phusion-Plus high-fidelity DNA polymerase and confirmed by whole plasmid sequencing (Plasmidsaurus, USA). The murine cDNAs for FERRY3 and TBCK were amplified from cDNA prepared from mouse kidney RNA. RNA was extracted from frozen kidney tissue using TRIZOL (Invitrogen). RNA integrity was analyzed by electrophoresis. Moloney murine leukemia virus reverse transcriptase (Invitrogen) was used to synthesize cDNA with random primers.

### Cell Experiments and Immunoblotting

HEK293 cells (ATCC® CRL-1573) were transiently transfected with the indicated expression constructs (see Table S1). Cells were maintained in DMEM (Gibco) supplemented with 10% fetal bovine serum (FBS) at 37°C in a humidified atmosphere containing 5% CO_2_. Transfections were performed at approximately 70–80% confluence using Lipofectamine 2000 (Life Technologies), following the manufacturer’s protocol.

Forty-eight hours post-transfection, cells were harvested and lysed in buffer containing 50 mM Tris-HCl (pH 7.5), 1 mM EGTA, 1 mM EDTA, 50 mM sodium fluoride, 5 mM sodium pyrophosphate, 1 mM sodium orthovanadate, 1% (w/v) Nonidet P-40, and 270 mM sucrose, supplemented with protease inhibitors (Complete tablets, Roche Applied Science) and 10 mM 1,10-phenanthroline. Protein concentrations were determined using a BCA assay (Bio-Rad), and subsequent immunoblot analyses were carried out as described above.

For live-cell imaging experiments, cells were plated onto glass-bottom culture dishes (Nest, 801002) and transfected as described. Fluorescently tagged constructs were used to enable visualization, and images were acquired between 24 and 48 hours after transfection.

### Immunoprecipitation

Recombinant FLAG-tagged proteins were immunoprecipitated using anti-FLAG M2 magnetic beads (Sigma) according to the manufacturer’s guidelines. In brief, 1 mg of total protein lysate was diluted in immunoprecipitation buffer composed of 1× TBS, 1% Triton X-100, 1 mM EDTA (pH 7.5), 1 mM sodium orthovanadate, 10 mM sodium pyrophosphate, and 50 mM sodium fluoride. Magnetic FLAG beads (30 µL) were added to the lysate and incubated for 2 h at 4°C with gentle agitation. Following incubation, beads were washed three times with 500 µL of TBS containing 0.05% Tween. Protein complexes were eluted using a denaturing buffer (2× Laemmli) supplemented with SDS, glycerol, bromophenol blue, and β-mercaptoethanol.

For HA-tagged recombinant proteins, immunoprecipitation was carried out using anti-HA magnetic beads (Pierce) following the supplier’s protocol. Briefly, 15 µL of beads were pre-washed three times with wash buffer (TBS supplemented with 0.1% Tween) prior to incubation with 1 mg of protein lysate. The mixture was incubated for 20 min at room temperature. Beads were subsequently washed three times and bound proteins were eluted in Laemmli sample buffer.

### Hypertonic Stimulation and Live-Cell Imaging

For hypertonic stimulation experiments, HEK293 cells were transiently transfected as described above and maintained under standard culture conditions. At the time of imaging, baseline images were first acquired under control conditions. Cells were then exposed to a hypertonic stimulus by addition of NaCl to a final concentration of 200 mM. Following stimulation, images were collected again to evaluate the cellular response before and after osmotic challenge.

For live-cell imaging, transfected cells were seeded onto glass-bottom dishes and visualized using fluorescently tagged constructs. Images were acquired immediately prior to stimulation and again after NaCl treatment under identical imaging settings.

### Structural homology-based search using the DALI server

Independent structural similarity searches were conducted with the DALI server ^27,28^ using four bait structures: the NRBP1 AlphaFold model, the SPAK AlphaFold model, the WNK2-CCT1 AlphaFold model, and the WNK1-CCT2 PDB structure (5G3Q). Searches targeted structures from the human AlphaFold Protein Structure Database. Candidate structures were retained for further analysis if they were identified by all four bait queries and had an average DALI Z-score of at least 4, a conservative threshold above the generally significant level (Z-score > 2). All aligned structures from each bait query were also manually inspected to verify fold similarity and exclude spurious matches.

### Structural homology-based search using Foldseek

We employed Foldseek ^29^ to identify structural homologs of the WNK kinase conserved C-terminal (CCT) domain. Eight structural “baits” were used to query the AFDB-proteome, including six experimental structures (PDB: 5G3Q, 2V3S, 6FBK, 7O86, 7OKW, 2LRU) and two AlphaFold2 models (NRBP1: AF-Q9UHY1; NRBP2: AF-Q91V36). To prioritize high-confidence candidates, raw hits were processed using a custom R pipeline. Data refinement was performed according to the following heuristic criteria.

Target descriptions were standardized to unify synonymous protein name and remove redundant descriptors. Then we retained only those targets with a Foldseek probability ≥0.1. To ensure fold robustness, candidates were required to be detected by at least four independent baits (n>3) and identified in two or more distinct species.

For each bait, only the top 50 targets by probability were included. Uncharacterized or generic “kinase domain-containing” proteins were excluded to focus on annotated functional contexts. The final filtered matrix was subjected to hierarchical clustering (Euclidean distance) and visualized using the heatmap package in R. This approach allowed for the identification of conserved structural clusters across the different CCT baits used in the study.

### Statistical Analysis

All statistical analyses were carried out using R software (version 2024.04.2+764). Comparisons between two independent groups were performed using an unpaired Student’s t-test. Data are expressed as mean ± standard deviation (SD), and a p-value < 0.05 was considered statistically significant.

For experiments involving more than two groups, statistical differences were assessed using one-way analysis of variance (ANOVA), followed by Tukey’s multiple comparisons test to identify group-specific differences.

## Acknowledgments

We thank Dario Alessi for the for insightful discussions and critical input and Marino Zerial for providing the plasmids encoding the human FERRY3 and TBCK proteins.

GMA is a doctoral student from the “Programa de Doctorado en Ciencias Biomédicas” of the “Universidad Nacional Autónoma de México” (UNAM) and received graduate student Fellowship CVU 942671 from “Secretaría de Ciencia, Humanidades, Tecnología e Innovación” (SECIHTI).

ERJ is a fellow in the General Office for Health Quality and Education, Ministry of Health (Dirección General de Calidad y Educación en Salud, Secretaría de Salud), Mexico.

MCB was supported by a KidneyCure Joseph V. Bonventre Career Development Grant, by SECIHTI grant CBF-2025-I-2028, and the 2025 Call to Support Health Science Research by the “Instituto Nacional de Ciencias Médicas y Nutrición Salvador Zubirán” (NMM-2147-25-31-1).

GG was supported by NIDDK R01 grant DK51496 and by the DGAPA-UNAM grant IN203025.

## Author contributions

Conceptualization: MCB, GMA, ERO; Methodology: GMA, ERO, MCB; Investigation: GMA, ERO, MLC, IDO, JB, HCC, ERJ, ROP, AMS, NV; Visualization: GMA, ERO, MCB; Supervision: MCB, GG, Writing: GMA, ERO, MCB.

## Data and materials availability

All data are available in the main text or the supplementary materials. Plasmids are available from MCB.

