## Supplementary material for "Determinants of CCT–motif specificity in WNK signaling and expansion of CCT-like domains"

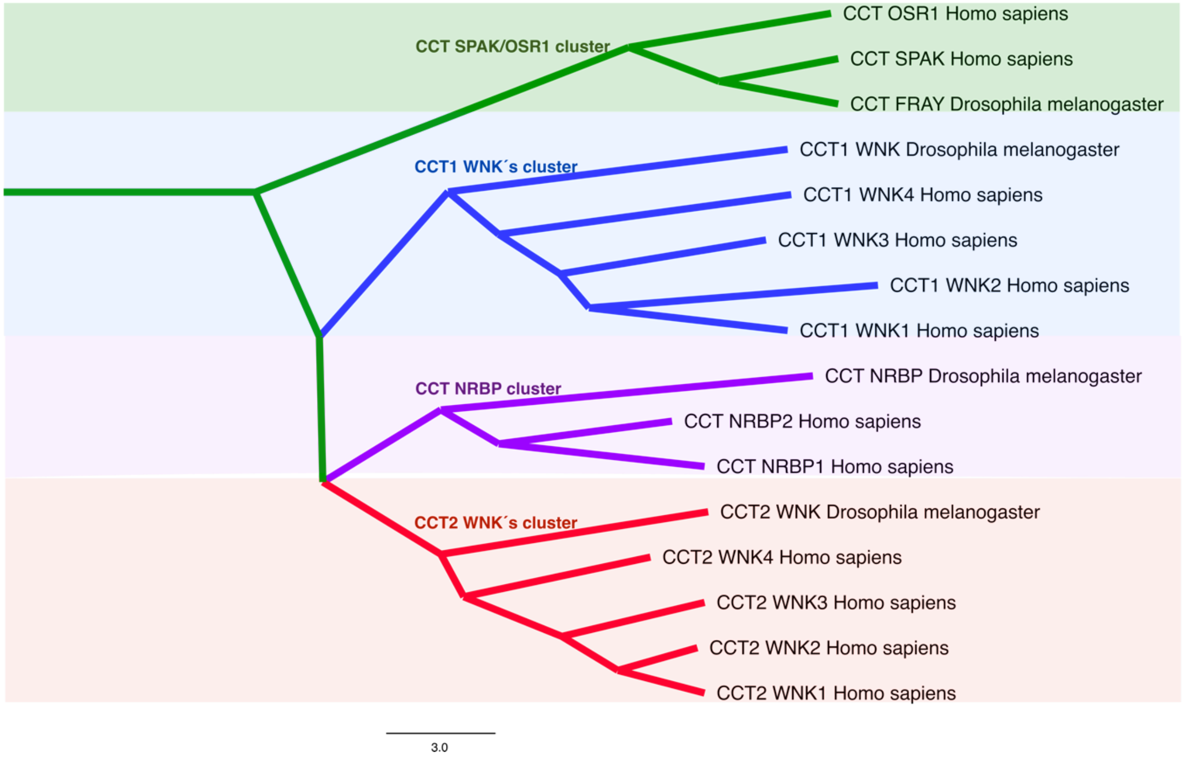


**Supplementary Figure 1. Structural tree of characterized CCT domains.** This tree was generated by structurally comparing AlphaFold models of CCT domains from humans and *Drosophila melanogaster* using the Dali server. Distinct structural clusters are represented by different colors: green for the SPAK/OSR1 cluster, blue for the WNK CCT1 cluster, purple for the NRBP CCT cluster, and red for the WNK CCT2 cluster.

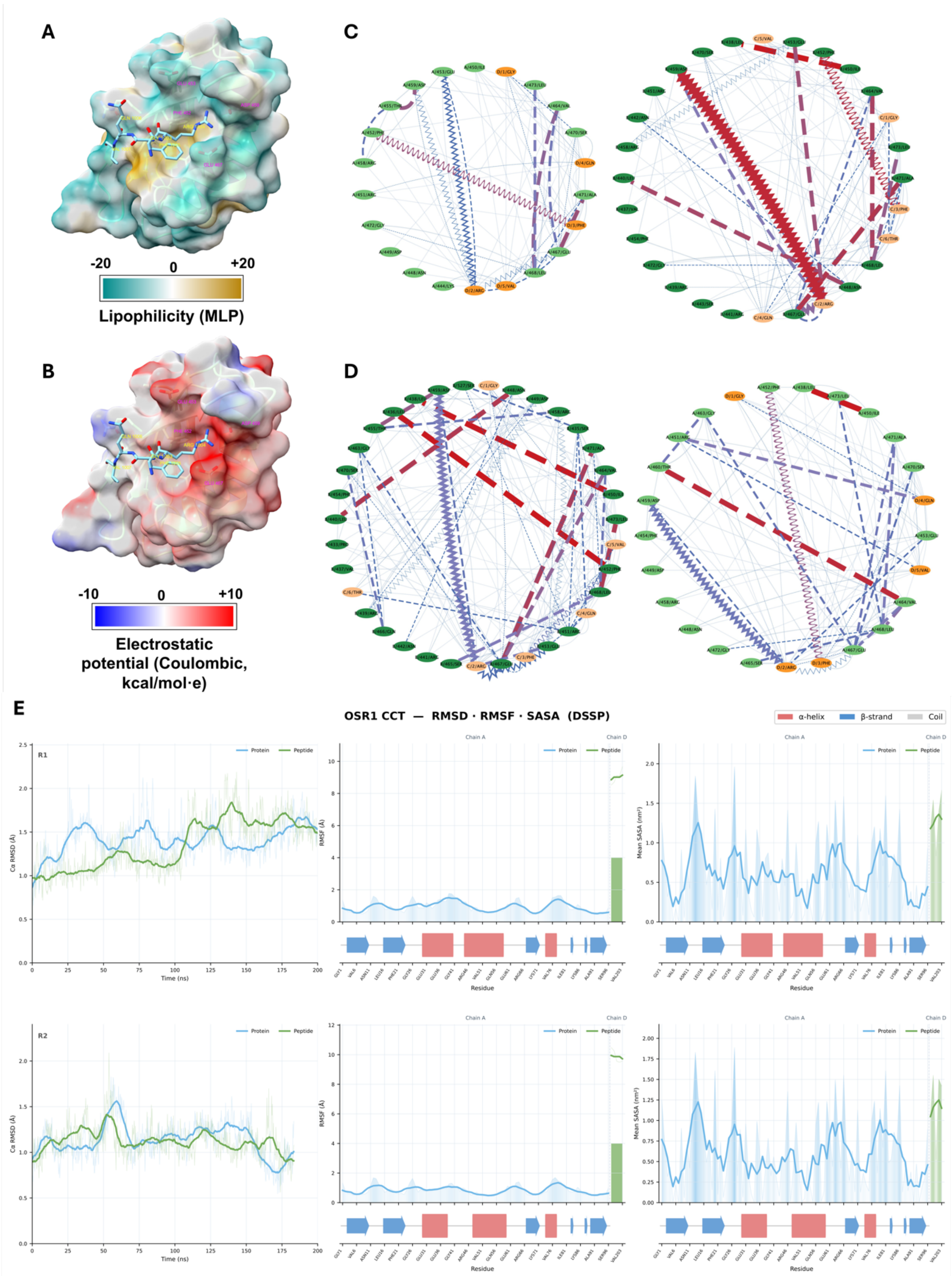


**Supplementary Figure 2. Structural analysis of the OSR1 CCT domain and its interaction network. (A-B)** Structure of the OSR1 CCT domain in complex with the WNK4 RFQV peptide (PDB ID: 2V3S). **(A)** Molecular lipophilicity potential (MLP) mapped onto the solvent-excluded surface (SES); the color gradient ranges from brown (hydrophobic) to blue (hydrophilic) based on the Ghose-Crippen atom-type scale. **(B)** Molecular surface colored by Coulombic electrostatic potential; red regions indicate negative potential (<−10 kcal/mol·e), blue regions indicate positive potential (>+10 kcal/mol·e), and white indicates neutral potential. Surfaces were rendered in ChimeraX v1.x using a probe radius of 1.4 Å. **(C-D)** Graph representation of the molecular interactions between the CCT domain and the RFQV peptide generated in Cytoscape. Panels show duplicate analysis of the dimer, highlighting the topology of intermolecular contacts. **(E)** Molecular Dynamics (MD) analysis of the OSR1 CCT-βϕ peptide crystallographic complex. Plots show RMSD (left), RMSF (center), and Mean SASA (right). Top panels represent replicate 1 and bottom panels represent replicate 2. The protein is shown in blue and the βϕ peptide in green. Secondary structure is indicated in the lower sections of the RMSF and SASA plots.

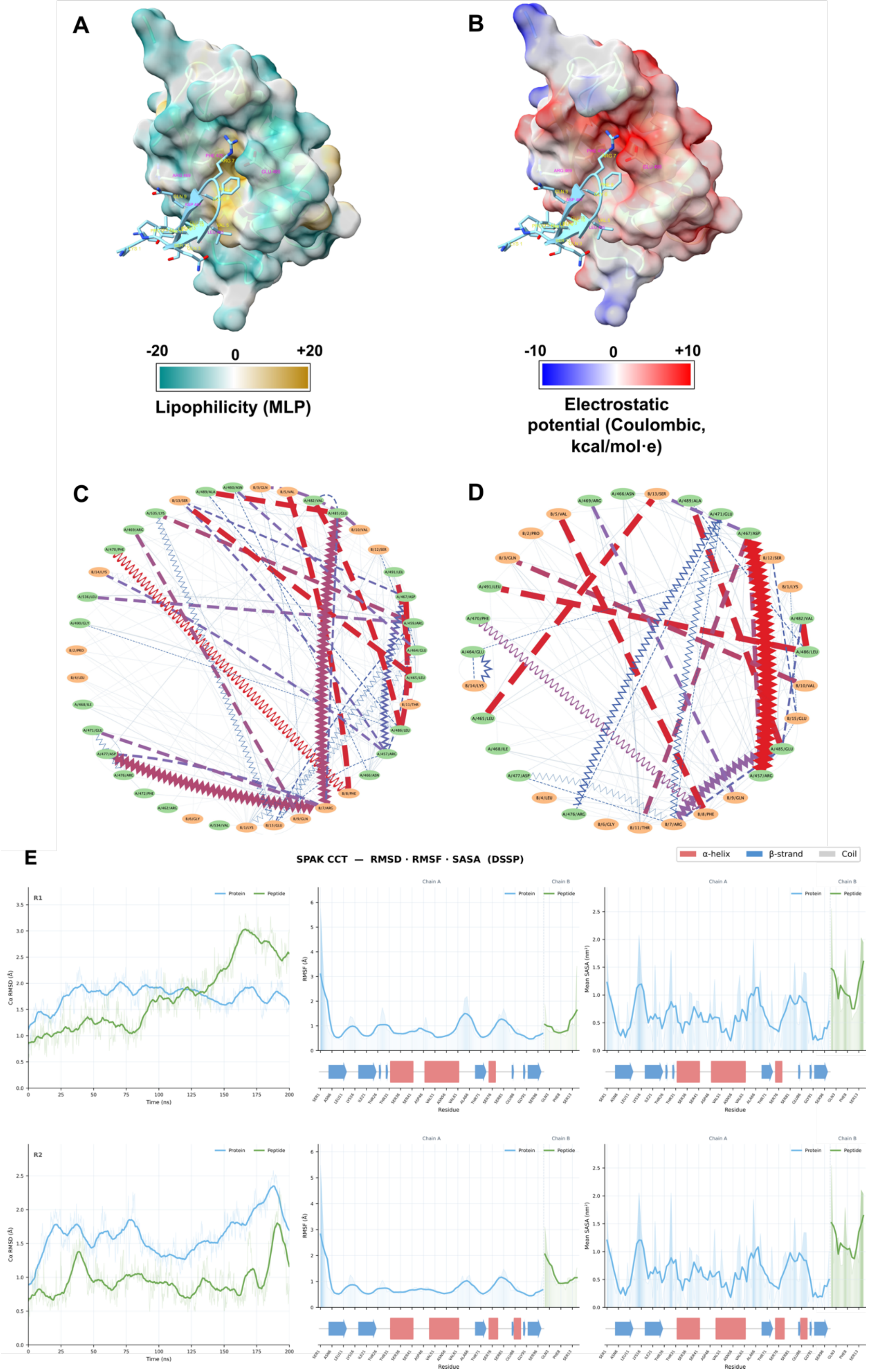
A
B
-20
0
+20
Lipophilicity (MLP)
-10
0
+10
Electrostatic potential (Coulombic, kcal/mol·e)
C
D
E
SPAK CCT — RMSD · RMSF · SASA (DSSP)
α-helix
β-strand
Coil
R1
R2
Protein
Peptide
Chain A
Chain B
Time (ns)
Residue

**Supplementary Figure 3. Structural modeling and interaction network of the SPAK CCT domain. (A-B)** Structural model of the SPAK CCT domain in complex with the extended WNK4 peptide (sequence: KPQLVGRFQVSSKE), obtained via AlphaFold 3. **(A)** MLP surface ranging from brown (hydrophobic) to blue (hydrophilic). **(B)** Molecular surface colored by Coulombic electrostatic potential (<−10 to >+10 kcal/mol·e). **(C-D)** Graph representation of the molecular interactions generated in Cytoscape. **(E)** MD analysis of the SPAK CCT-βϕ peptide from the AlphaFold model. Plots show RMSD (left), RMSF (center), and Mean SASA (right). Top panels represent replicate 1 and bottom panels represent replicate 2. Protein in blue, βϕ peptide in green. Secondary structure is depicted in the lower parts of the RMSF and SASA plots.

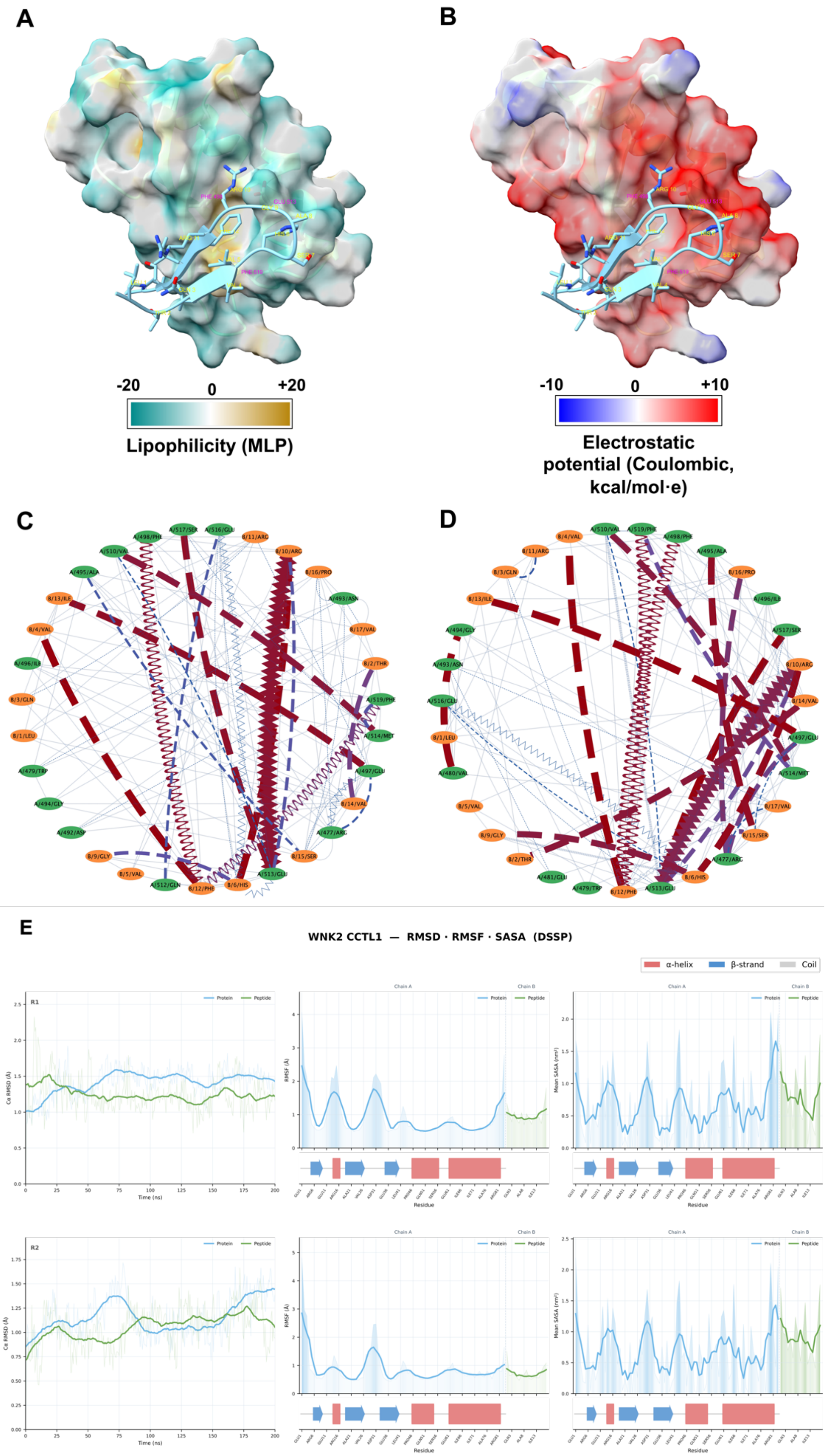

A
B
-20
0
+20
Lipophilicity (MLP)
-10
0
+10
Electrostatic potential (Coulombic, kcal/mol·e)
C
D
E
WNK2 CCTL1 — RMSD · RMSF · SASA (DSSP)
α-helix
β-strand
Coil
R1
R2
Protein
Peptide
Chain A
Chain B
Time (ns)
Residue

**Supplementary Figure 4. Structural analysis of the human WNK2 CCT-like 1 domain. (A-B)** Crystal structure of the human WNK2 CCT-like 1 domain in complex with the WNK1 RFXV peptide (PDB ID: 3FV0). **(A)** MLP surface. **(B)**Coulombic electrostatic potential surface (<−10 to >+10 kcal/mol·e). **(C-D)** Graph representation of the molecular interactions generated in Cytoscape. **(E)** MD analysis of the WNK2 CCT1 domain based on the crystallographic complex PDB: 6FBK. Plots show RMSD (left), RMSF (center), and Mean SASA (right). Upper panels represent replicate 1 and lower panels represent replicate 2. Protein in blue, βϕ peptide in green. Secondary structure is shown in the lower sections of the RMSF and SASA plots.

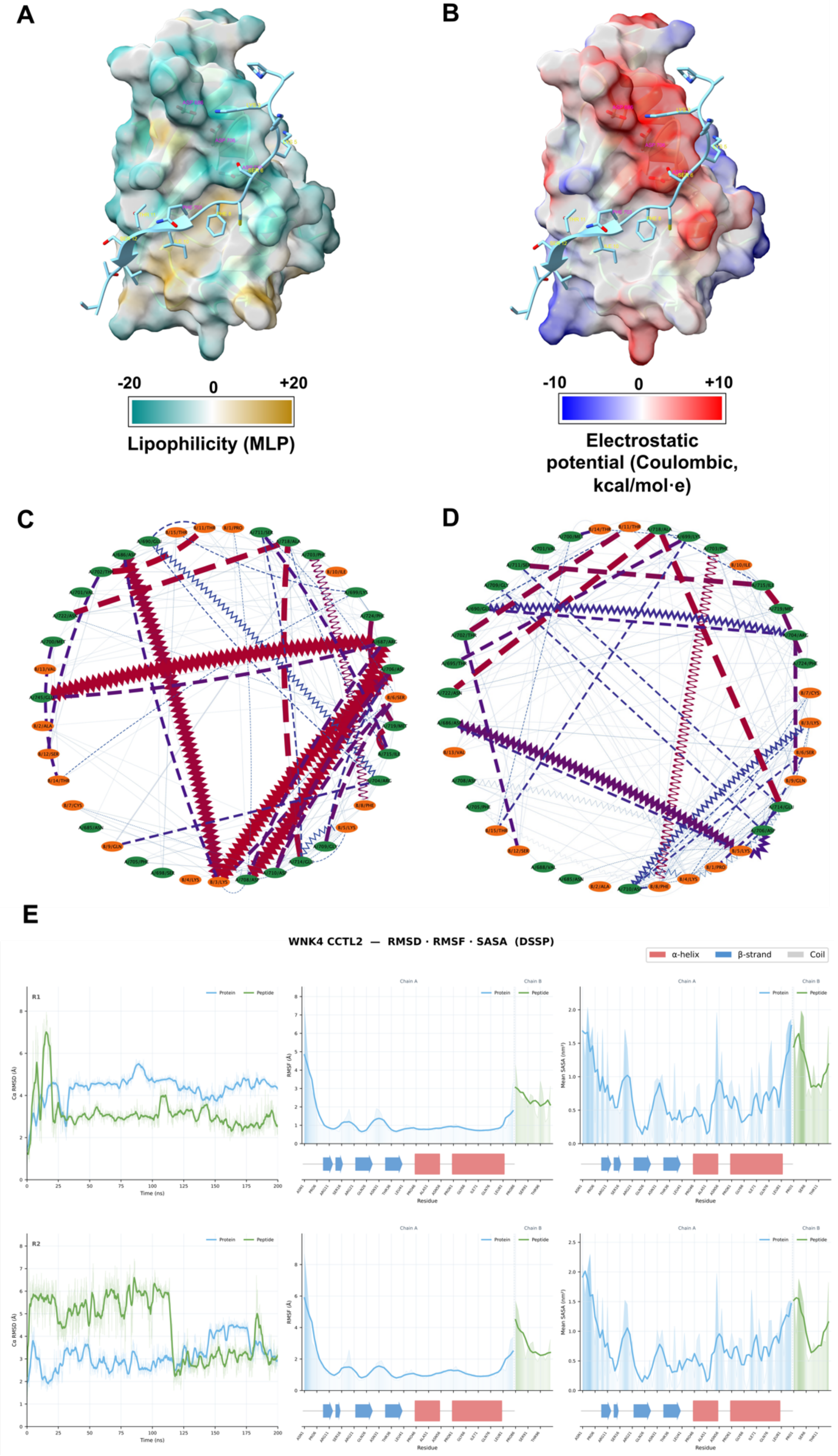
A
B
-20
0
+20
Lipophilicity (MLP)
-10
0
+10
Electrostatic potential (Coulombic, kcal/mol·e)
C
D
E
WNK4 CCTL2 — RMSD · RMSF · SASA (DSSP)
α-helix
β-strand
Coil
R1
R2
Protein
Peptide
Chain A
Chain B
Time (ns)
Residue

**Supplementary Figure 5. Structural modeling of the WNK4 CCT2 domain and its interaction with TSC22D2. (A-B)** Structural model of the WNK4 CCT2 domain in complex with the TSC22D2 βϕ A peptide (sequence: PAKKKSCFQITSVTT), obtained via AlphaFold 3. **(A)** MLP surface. **(B)** Coulombic electrostatic potential surface (<−10 to >+10 kcal/mol·e). **(C-D)** Graph representation of the molecular interactions generated in Cytoscape. **(E)** MD analysis of the WNK4 CCT2-βϕ based on the AlphaFold model. Plots show RMSD (left), RMSF (center), and Mean SASA (right). Upper panels show replicate 1 and bottom panels show replicate 2. Protein in blue, βϕ peptide in green. Secondary structure is indicated in the lower sections of the RMSF and SASA plots.

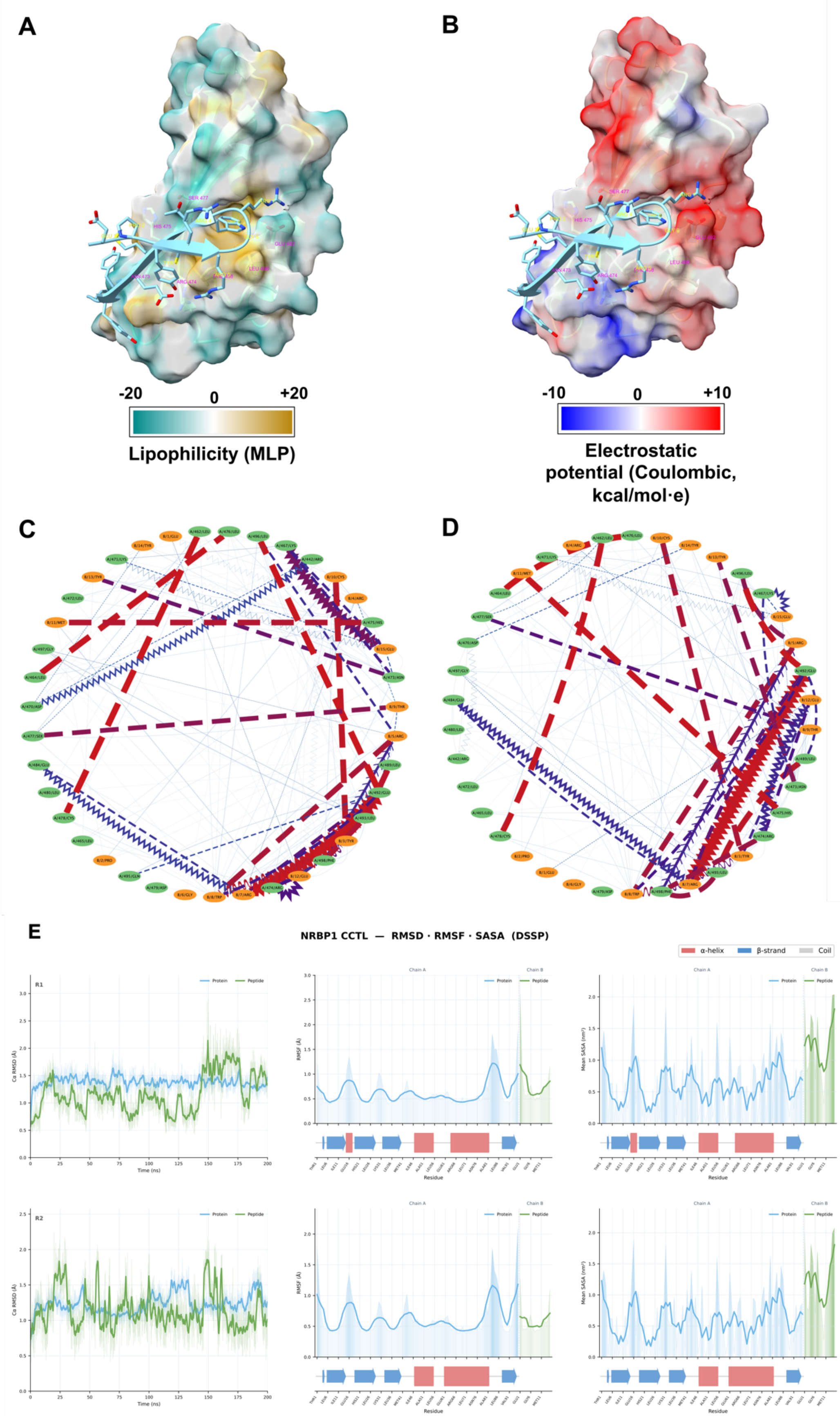
A
B
-20
0
+20
Lipophilicity (MLP)
-10
0
+10
Electrostatic potential (Coulombic, kcal/mol·e)
C
D
E
NRBP1 CCTL — RMSD · RMSF · SASA (DSSP)
α-helix
β-strand
Coil

**Supplementary Figure 6. Structural modeling of the NRBP1 CCT domain and its interaction with TSC22D2. (A-B)**Structural model of the NRBP1 CCT domain in complex with the TSC22D2 βϕ C peptide (sequence: EPYRRGRWTCMEYYE), obtained via AlphaFold 3. **(A)** MLP surface. **(B)** Coulombic electrostatic potential surface ($<-10$ to $>+10$ kcal/mol·e). **(C-D)** Graph representation of the molecular interactions generated in Cytoscape. **(E)** MD analysis of the NRBP1 CCT-βϕ derived from the AlphaFold model. Plots show RMSD (left), RMSF (center), and Mean SASA (right). Top panels represent replicate 1 and bottom panels represent replicate 2. Protein in blue, βϕ peptide in green. Secondary structure is depicted in the lower sections of the RMSF and SASA plots.

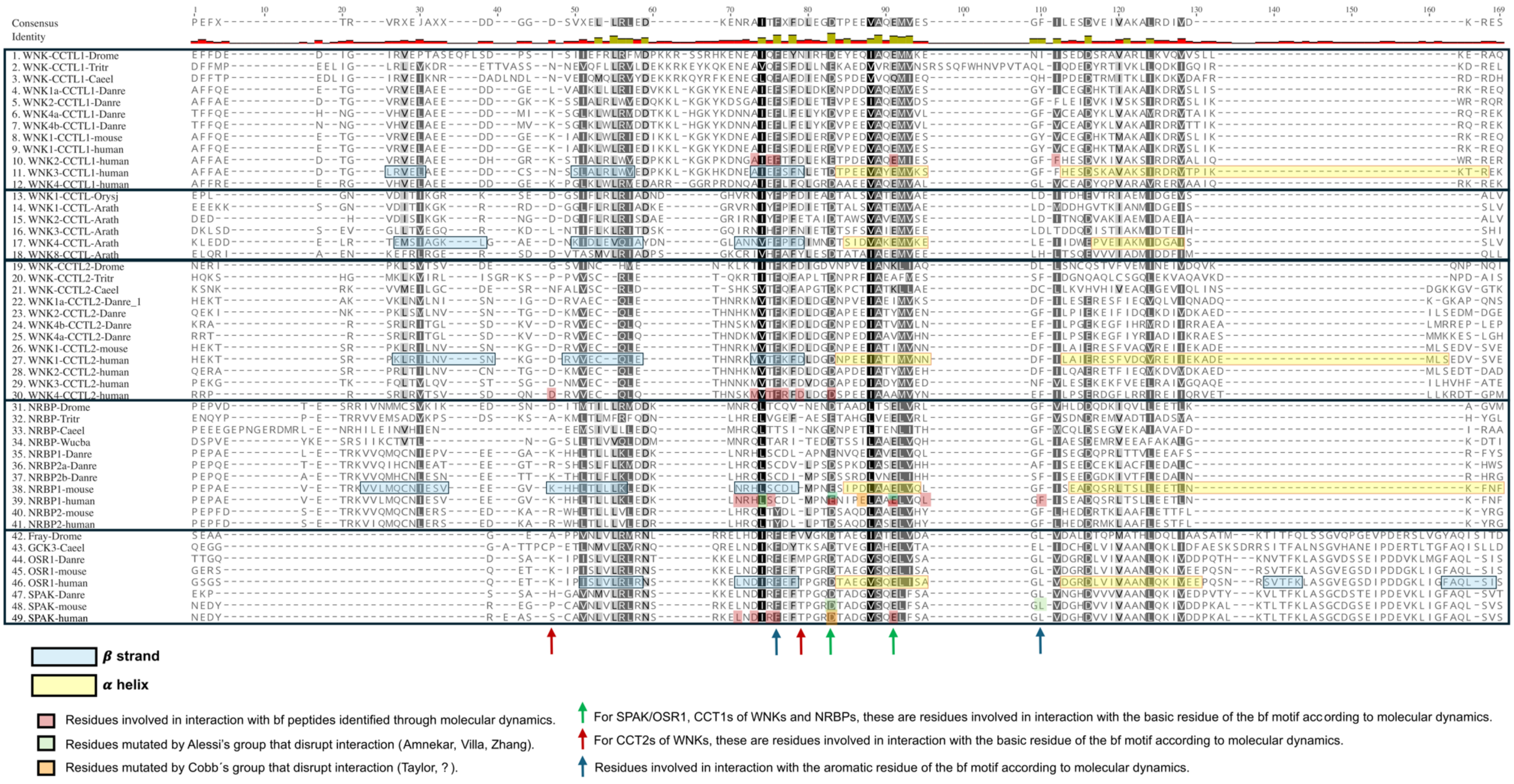
Consensus
Identity
1. WNK-CCTL1-Drome
2. WNK-CCTL1-Tritr
3. WNK-CCTL1-Caeel
4. WNK1a-CCTL1-Danre
5. WNK2-CCTL1-Danre
6. WNK4a-CCTL1-Danre
7. WNK4b-CCTL1-Danre
8. WNK1-CCTL1-mouse
9. WNK1-CCTL1-human
10. WNK2-CCTL1-human
11. WNK3-CCTL1-human
12. WNK4-CCTL1-human
13. WNK1-CCTL-Orysj
14. WNK1-CCTL-Arath
15. WNK2-CCTL-Arath
16. WNK3-CCTL-Arath
17. WNK4-CCTL-Arath
18. WNK8-CCTL-Arath
19. WNK-CCTL2-Drome
20. WNK-CCTL2-Tritr
21. WNK-CCTL2-Caeel
22. WNK1a-CCTL2-Danre_1
23. WNK2-CCTL2-Danre
24. WNK4b-CCTL2-Danre
25. WNK4a-CCTL2-Danre
26. WNK1-CCTL2-mouse
27. WNK1-CCTL2-human
28. WNK2-CCTL2-human
29. WNK3-CCTL2-human
30. WNK4-CCTL2-human
31. NRBP-Drome
32. NRBP-Tritr
33. NRBP-Caeel
34. NRBP-Wucba
35. NRBP1-Danre
36. NRBP2a-Danre
37. NRBP2b-Danre
38. NRBP1-mouse
39. NRBP1-human
40. NRBP2-mouse
41. NRBP2-human
42. Fray-Drome
43. GCK3-Caeel
44. OSR1-Danre
45. OSR1-mouse
46. OSR1-human
47. SPAK-Danre
48. SPAK-mouse
49. SPAK-human
β strand
α helix
Residues involved in interaction with bf peptides identified through molecular dynamics.
Residues mutated by Alessi's group that disrupt interaction (Amnekar, Villa, Zhang).
Residues mutated by Cobb´s group that disrupt interaction (Taylor, ? ).
For SPAK/OSR1, CCT1s of WNKs and NRBPs, these are residues involved in interaction with the basic residue of the bf motif according to molecular dynamics.
For CCT2s of WNKs, these are residues involved in interaction with the basic residue of the bf motif according to molecular dynamics.
Residues involved in interaction with the aromatic residue of the bf motif according to molecular dynamics.

**Supplementary Figure 7. Multiple sequence alignment of CCT domains across species.** Alignment of representative CCT domains indicating secondary structure elements and key residues for βϕ motif interaction. **Secondary structure:** β-strands are boxed in sky blue and α-helices in yellow. **Functional residues:** Light red highlights indicate residues involved in βϕ peptide interaction identified through molecular dynamics; light green indicates residues whose mutation disrupts interaction (Alessi group); light orange indicates residues mutated by the Cobb group (Taylor et al., 2018). **Specific interaction markers: Green arrows:** Residues interacting with the basic residue of the βϕ motif in SPAK/OSR1, WNK CCT1s, and NRBPs. **Red arrows:** Residues interacting with the basic residue of the βϕ motif specifically in WNK CCT2 domains. **Blue arrows:** Residues involved in interaction with the aromatic residue of the βϕ motif.

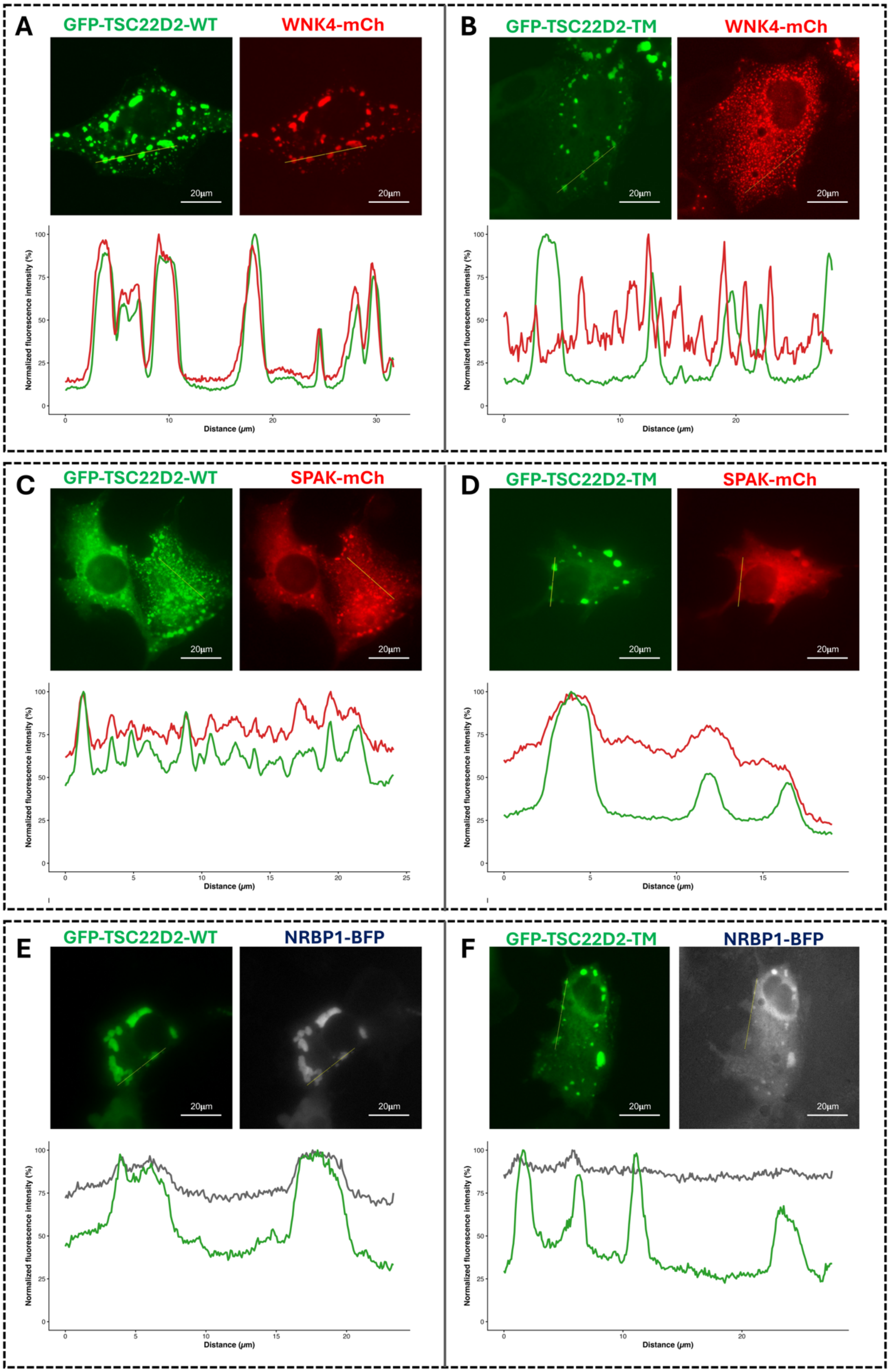

A
GFP-TSC22D2-WT
WNK4-mCh
20µm
20µm
Normalized fluorescence intensity (%)
Distance (µm)
B
GFP-TSC22D2-TM
WNK4-mCh
20µm
20µm
Normalized fluorescence intensity (%)
Distance (µm)
C
GFP-TSC22D2-WT
SPAK-mCh
20µm
20µm
Normalized fluorescence intensity (%)
Distance (µm)
D
GFP-TSC22D2-TM
SPAK-mCh
20µm
20µm
Normalized fluorescence intensity (%)
Distance (µm)
E
GFP-TSC22D2-WT
NRBP1-BFP
20µm
20µm
Normalized fluorescence intensity (%)
Distance (µm)
F
GFP-TSC22D2-TM
NRBP1-BFP
20µm
20µm
Normalized fluorescence intensity (%)
Distance (µm)

**Supplementary Figure 8. Co-localization analysis of TSC22D2 with WNK4, SPAK, and NRBP1.** Evaluation of intracellular interaction via fluorescence microscopy and line intensity profiles. **(A-B)** Co-expression of WNK4-mCherry with TSC22D2-GFP-WT **(A)** or the triple mutant (TM) variant **(B)**. **(C-D)** Co-expression of SPAK-mCherry with TSC22D2-GFP-WT **(C)** or the TM variant **(D)**. **(E-F)** Co-expression of NRBP1-BFP with TSC22D2-GFP-WT **(E)** or the TM variant **(F)**. Each panel displays individual fluorescence channels and a line intensity profile plot generated in FIJI across a representative ROI. Profiles allow for comparison of spatial distribution of fluorescence peaks.

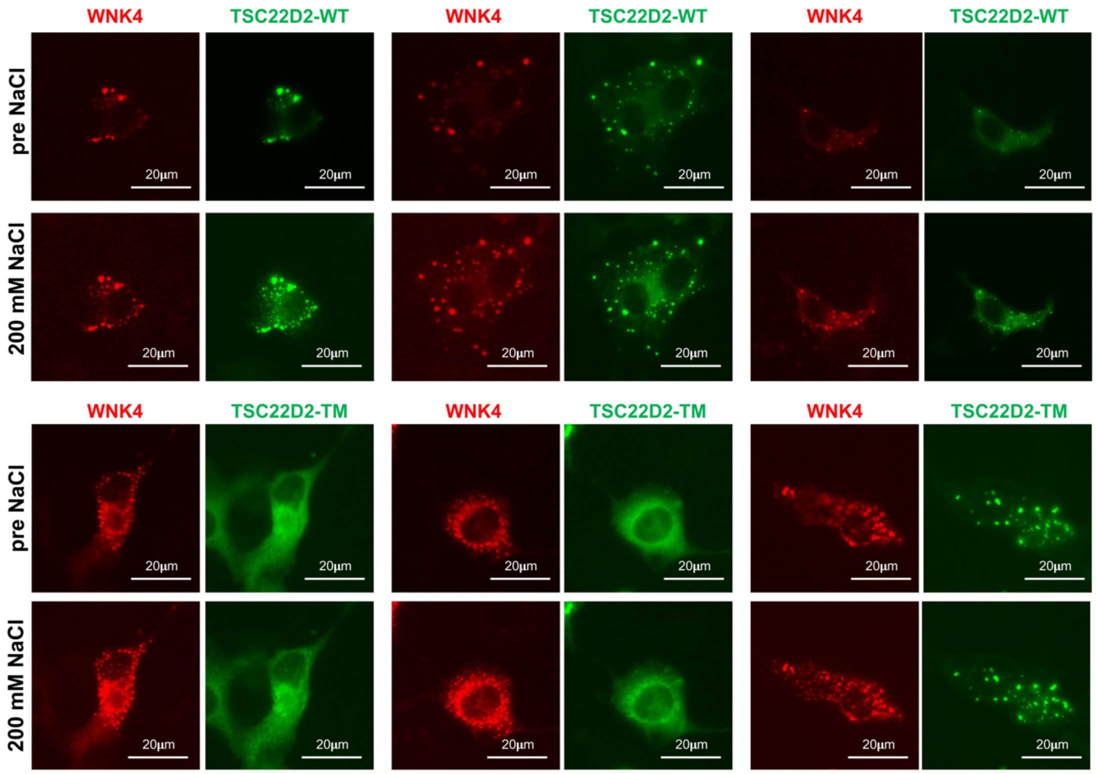


**Supplementary Figure 9. TSC22D2 recruitment dynamics in response to hypertonic stress.** Live-cell fluorescence microscopy of cells co-transfected with WNK4-mCherry and TSC22D2-GFP variants. **(A-C)** Representative fields of cells expressing TSC22D2-WT and WNK4-mCherry under basal conditions and after the addition of 200 mM NaCl. **(D-F)** Representative fields of the TSC22D2-GFP triple βϕ mutant. Unlike the WT variant, the triple mutant shows no increase in cytoplasmic particle formation following hypertonic stress, suggesting these motifs are essential for the osmotic stress response.

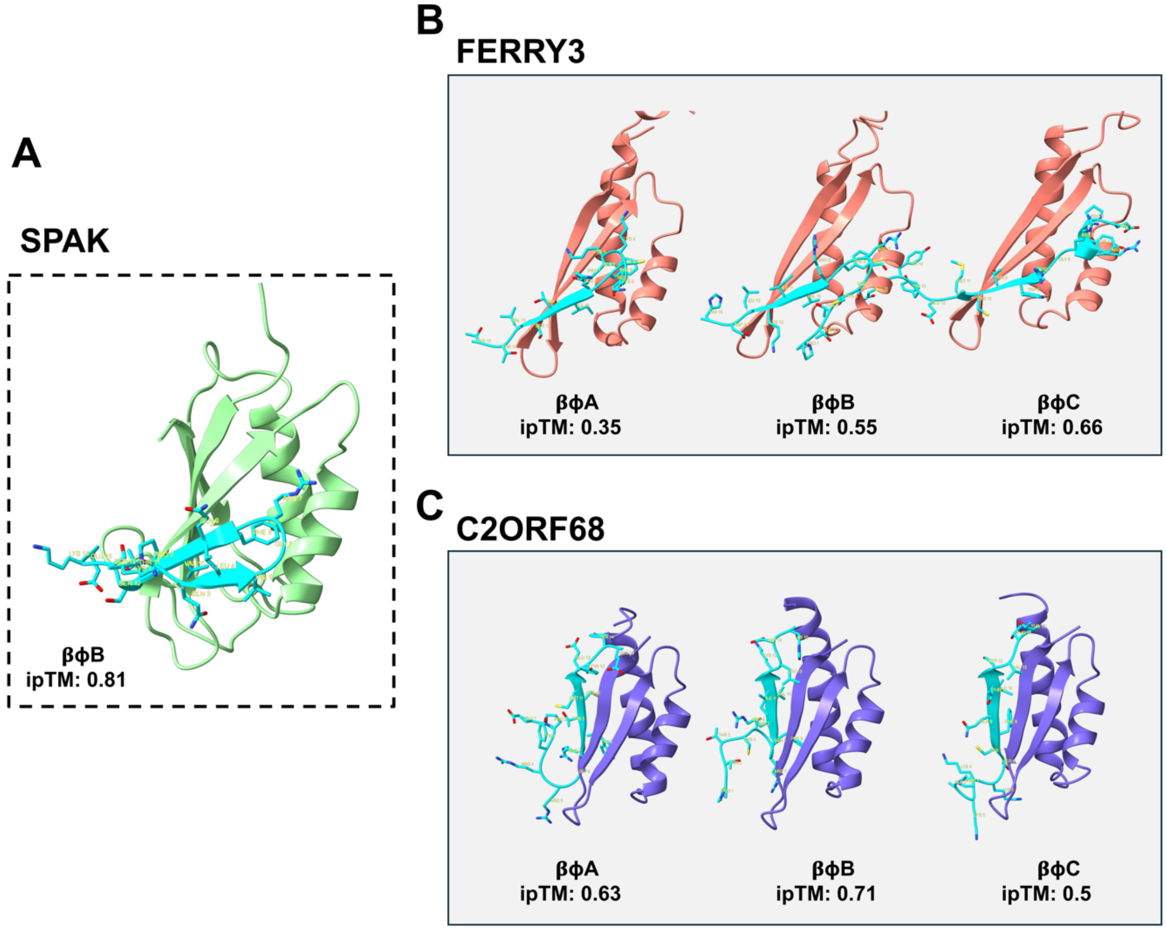


**Supplementary Figure 10 AF3 structural modeling of βϕ-motif interactions of FERRY3 and C2ORF68 putative CCT domains with known βϕ motifs.**

**A)** AlphaFold3 (AF3) prediction of the human SPAK CCT domain (pale green) in complex with the TSC22D2 βϕ-motif B (cyan). The AF3 calculated ipTM value is shown below. The model illustrates the conserved binding pocket of the CCT domain coordinating the hydrophobic residues of the βϕ motif. **B)** Structural model of the FERRY3 putative CCT domain (salmon) in complex with the βϕ motifs A, B, and C from TSC22D2 (cyan). The AF3 calculated ipTM values are shown below. The model reveals an interaction architecture for ferry3 that is analogous to the SPAK complex, highlighting a conserved mechanism for the recognition of multiple βϕ linear motifs within a single target protein. **C)** AF3-predicted complex of C2orf68 (purple) with the βϕ motifs A, B, and C of TSC22D2 (cyan). The AF3 calculated ipTM values are shown below.

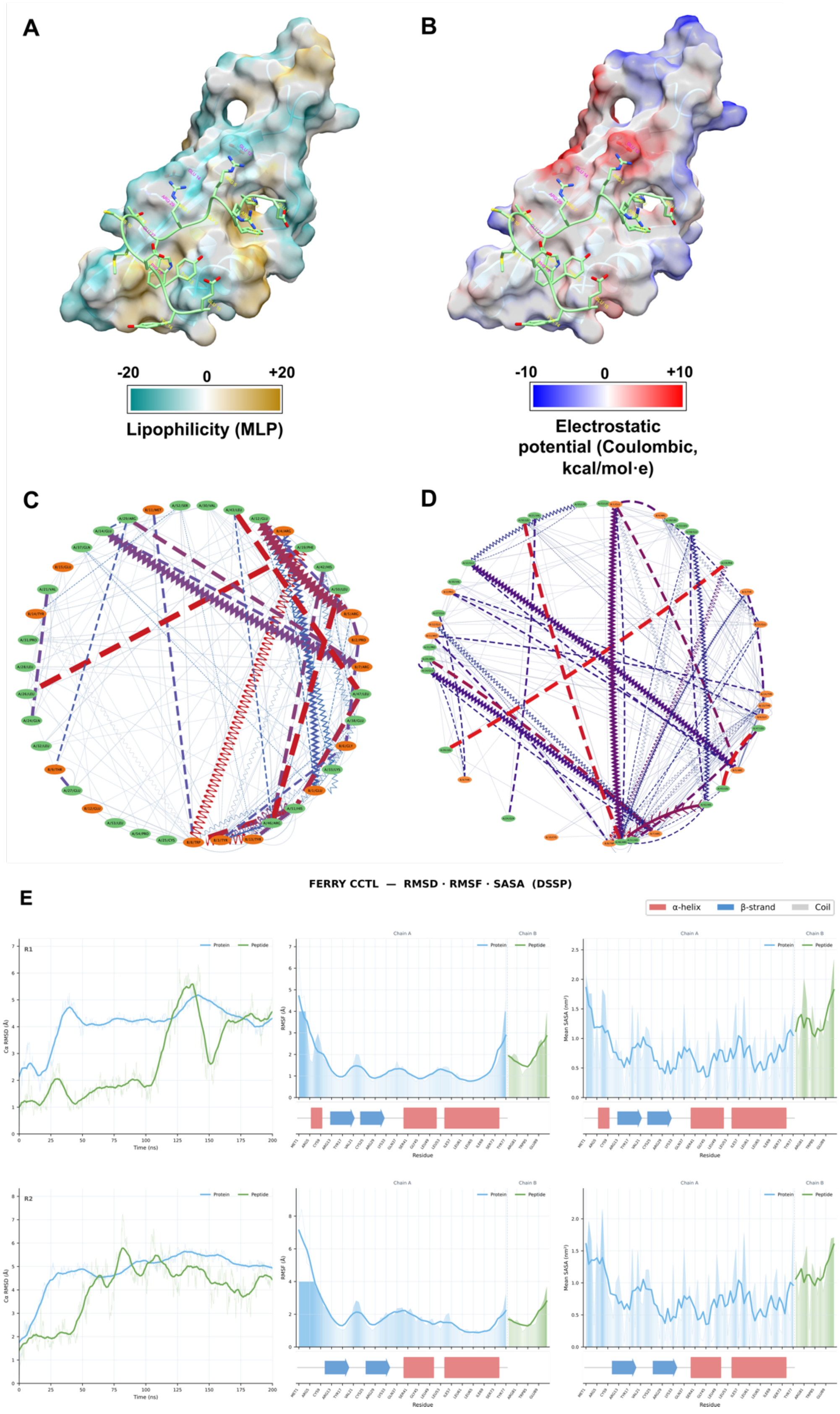
A
B
-20
0
+20
Lipophilicity (MLP)
-10
0
+10
Electrostatic potential (Coulombic, kcal/mol·e)
C
D
E
FERRY CCTL — RMSD · RMSF · SASA (DSSP)
α-helix
β-strand
Coil
R1
R2
Protein
Peptide
Chain A
Chain B
Cα RMSD (Å)
Time (ns)
RMSF (Å)
Mean SASA (nm²)
Residue

**Supplementary Figure 11. Structural modeling of the FERRY3 CCT-like domain and its interaction with TSC22D2. (A-B)** Structural model of the FERRY3 CCT-like domain in complex with the TSC22D2 βϕ C peptide (sequence: EPYRRGRWTCMEYYE), obtained via AF3. **(A)** MLP surface. **(B)** Coulombic electrostatic potential surface (<−10 to >+10 kcal/mol·e). **(C-D)** Graph representation of molecular interactions generated in Cytoscape. **(E)** MD analysis of the FERRY3 CCT-βϕ based on the AlphaFold model. Plots show RMSD (left), RMSF (center), and Mean SASA (right). Top panels represent replicate 1 and bottom panels represent replicate 2. Protein in blue, βϕ peptide in green. Secondary structure is shown in the lower sections of the RMSF and SASA plots.

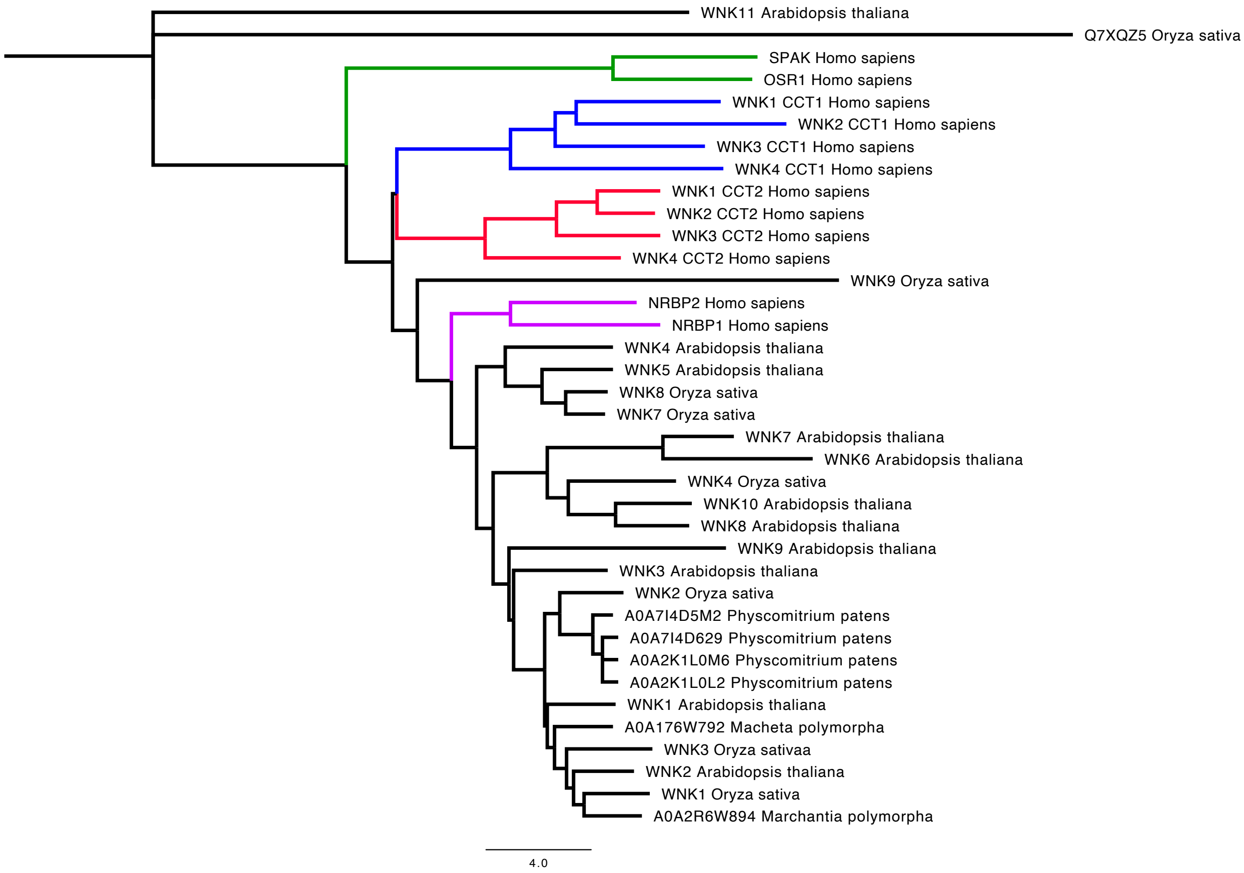


**Supplementary Figure 12. Structure-based clustering tree of putative CCT domains from plant WNK kinases.** Human reference clusters are shown in green for SPAK/OSR1 CCT, blue for WNK CCT1, red for WNK CCT2, and purple for NRBP. All plant serine/threonine kinases from Arabidopsis thaliana, Oryza sativa, Marchantia polymorpha, and Physcomitrium patens with putative CCT domains annotated in UniProt were included. Most putative plant CCT domains group within the human NRBP cluster.
